# Rewired meristem regulation drives morphological evolution of the one-leaf developmental system in *Monophyllaea glabra*

**DOI:** 10.64898/2026.08.05.742951

**Authors:** Shunji Nakamura, Ayaka Kinoshita, Hiroyuki Koga, Hirokazu Tsukaya

**Affiliations:** Department of Biological Sciences, Graduate School of Science, The University of Tokyo, Tokyo, 113-0033, Japan

## Abstract

Plants have evolved unconventional shoot systems with a “fuzzy” character that blurs organ boundaries, but the developmental programs driving this morphological diversity remain a mystery. One-leaf plants of the genus *Monophyllaea* are a striking example of this diversity: they develop a single indeterminate cotyledon instead of a conventional shoot system to form a unique structural unit termed a phyllomorph. Here, we integrate tissue-specific and single-nucleus transcriptomics with spatial gene expression analyses to characterize the meristems underlying this developmental system. The *Monophyllaea*-specific meristem of the phyllomorph retains a conserved shoot apical meristem-like transcriptional core but has a leaf lamina program superimposed. This chimeric transcriptional state integrates normally distinct developmental programs alongside altered phytohormone regulation. The one-leaf morphology emerges through the rewiring of conserved meristem regulatory networks rather than the passive suppression of meristem activity. The findings highlight the remarkable plasticity of developmental modules in plant morphological evolution.

## Introduction

Morphological diversification is a fundamental driver of evolution, arising from the modification of developmental programs that organize distinct cell types. Among the most extreme examples of such modification are lineages that deviate from canonical body plans^1,2^: the genus *Monophyllaea* (Gesneriaceae) exemplifies this by producing no stems or foliage leaves during the vegetative phase due to limited organogenic activity (**Figure 1A**)^3^. Instead, it establishes a unique one-leaf developmental system based on a structural unit known as the phyllomorph^4^. This developmental unit is composed of an indeterminately growing lamina and a petiolode, endowing the organ with both stem- and petiole-like characteristics.

**Figure 1.**
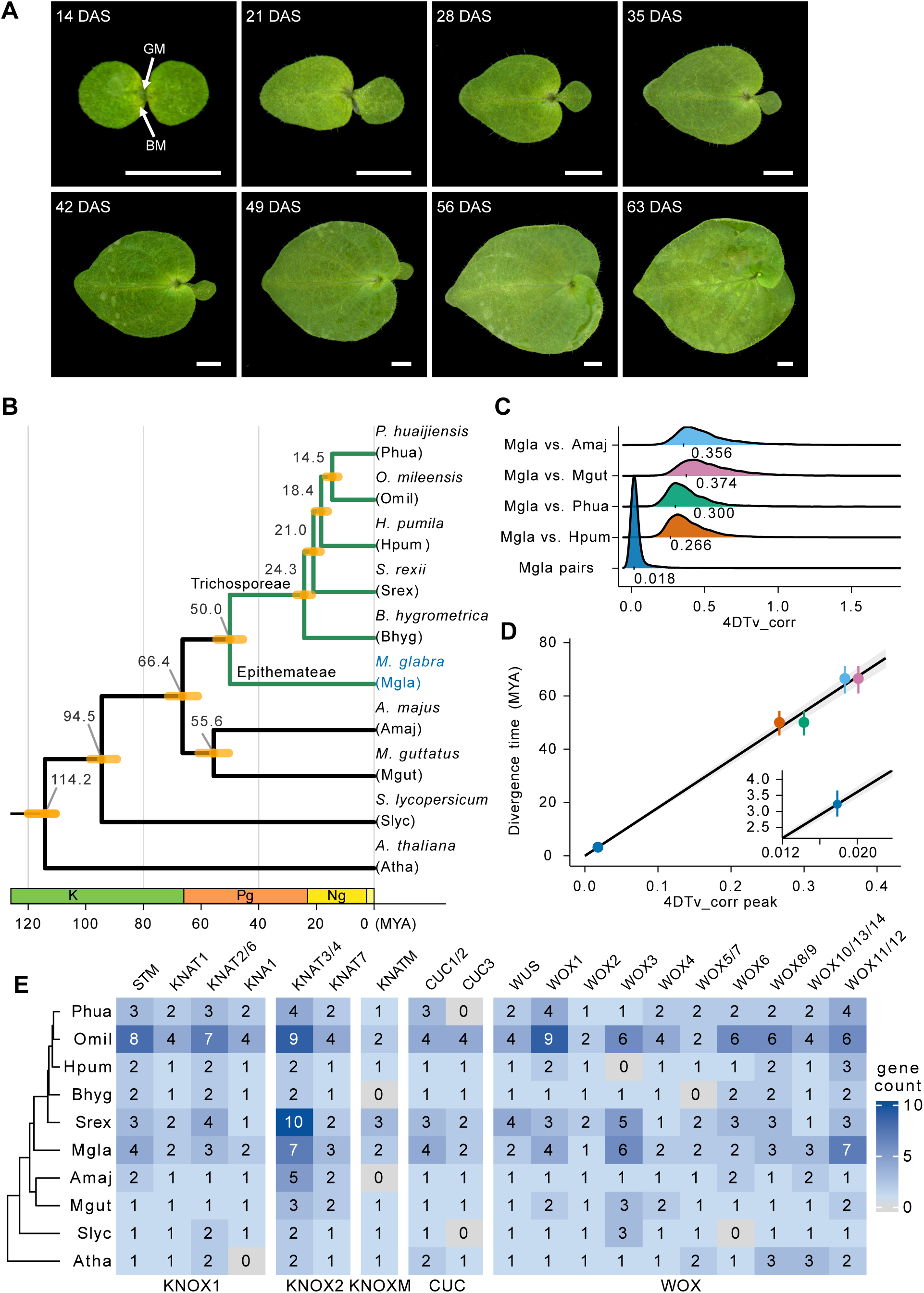
Unique developmental process and genomic evolution of *Monophyllaea glabra*. (A) Representative images of the characteristic developmental stages of *M. glabra* under short-day conditions. DAS, days after sowing; GM, groove meristem; BM, basal meristem. Scale bars = 2 mm. (B) Phylogenetic tree and estimated divergence times. Tree topology and node ages were inferred using 1727 single-copy orthologous groups, with divergence times estimated using MCMCtree with fossil/time calibrations. K, Cretaceous; Pg, Paleogene; Ng, Neogene. (C) Distributions of four-fold degenerate transversion (4DTv) distances. The plot displays 4DTv values for strict 1:1 interspecific orthologous pairs among selected species, as well as within-genome duplicated gene pairs of *M. glabra*. (D) Correlation between representative 4DTv peaks and estimated divergence times. Interspecific 4DTv peaks were plotted against MCMCtree-derived posterior node ages to fit a through-origin model. (E) Heatmap showing the estimated number of genes related to core meristem regulators (KNOX1, KNOX2, KNOXM, CUC, and WOX) across the analyzed species. Subcategories are labeled using *A. thaliana* gene names, except for KNA1, which is absent from *A. thaliana* and is therefore shown as KNOX1.

In angiosperms, the shoot apical meristem (SAM) serves as the central source of organization, maintaining a population of undifferentiated stem cells that sequentially produce above-ground organs^5–7^. While the SAM is highly conserved across angiosperms, certain lineages have evolved divergent architectures through its modification or loss^2^. *Monophyllaea* lack a conventional shoot system, and this architectural divergence is mediated by a groove meristem (GM) that functionally replaces the SAM^4,8,9^. This fundamental structural modification is particularly striking given that Gesneriaceae (∼150 genera, 3900 species) is a morphologically diverse pantropical family that largely maintains standard shoot architectures^10,11^. Understanding how the distinct architecture of *Monophyllaea* emerges from the conserved genetic toolkits shared with its conventional relatives remains a central question in evolutionary developmental biology^2,12–14^.

*Monophyllaea* members also possess a characteristic basal meristem (BM). Following germination, seedlings initially expand equal-sized cotyledons (isocotyly), before transitioning to an anisocotylous stage (**Figure 1A**). Notably, this asymmetric growth is governed by the BM without direct cell supply from the GM^15^. The BM forms specifically at the base of one cotyledon, termed the macrocotyledon, contributing to the indeterminate growth of the lamina, while growth of the opposing microcotyledon is suppressed. Concurrently, a petiolode meristem located beneath the GM contributes to thickening of the midrib. Upon transition to the reproductive phase, cell division in the GM is activated to form inflorescence and floral meristems capable of conventional organogenesis^3,16^. The phyllomorph of *Monophyllaea* is therefore considered an example of the enigmatic evolution of “fuzzy” morphology in which organ boundaries blur beyond traditional structural categories such as stems, leaves, and roots^1,3,17^. Similar morphological anomalies include the leaf-like shoot system of *Asparagus*, the thallus-like root system of Podostemaceae, and the bladder-bearing shoot of *Utricularia*^18–20^.

In model angiosperms such as *Arabidopsis thaliana*, SAM maintenance relies on a tightly regulated gene network that prevents differentiation. Class I KNOTTED 1-LIKE HOMEOBOX (KNOX1) family genes, such as *SHOOT MERISTEMLESS* (*STM*), play a pivotal role in this process by creating a high-cytokinin (CK)/low-gibberellin (GA) environment^21–23^. STM activates CK biosynthesis genes such as *ISOPENTENYLTRANSFERASE 7* (*IPT7*), and represses GA biosynthesis genes such as *GA20-oxidase* (*GA20ox*). This low-GA state is further reinforced by upregulation of the catabolic gene *GA2-oxidase* (*GA2ox*), a conserved regulatory target of KNOX1 transcription factors, thereby inhibiting differentiation in maize^24^. This hormonal balance functions in concert with the *WUSCHEL* (*WUS*)-*CLAVATA 3* (*CLV3*) feedback loop, which spatially defines the stem cell niche^25^.

However, unlike a conventional dome-shaped SAM, the *Monophyllaea* GM is groove-shaped and exhibits low organogenic activity. This raises the question of whether the GM operates through the same regulatory framework as a conventional SAM. To address this question, we selected *Monophyllaea glabra*, a rare annual species particularly suitable for molecular and developmental studies^26,27^. Our previous expression analyses suggested that the GM has a complex molecular identity. The GM expresses *STM* orthologs (*MgSTM*), supporting homology between the GM and the SAM^3,16,28^. The GM also expresses the ortholog of *ANGUSTIFOLIA 3*/*Arabidopsis thaliana GRF-*

*INTERACTING FACTOR 1* (*MgAN3*), which serves not only as a marker of leaf primordia, but also as a regulator of leaf meristem activity, suggesting that the GM has both SAM- and leaf-associated properties^3,29^. Furthermore, the expression domains of *MgSTM* and *CUP-SHAPED COTYLEDON* orthologs (*MgCUCs*) overlap within the GM, a pattern reminiscent of early embryogenesis rather than of a mature SAM^16,30^. However, these observations were based on a limited number of marker genes, and the broader molecular organization of the phyllomorph remains unresolved.

Here, we established a multi-omics platform for *M. glabra*, integrating a *de novo* draft genome assembly, tissue-specific bulk RNA sequencing (RNA-seq), single-nucleus RNA-seq, and spatial gene expression analyses. We demonstrate that the GM retains a conserved SAM-like transcriptional core, including the *STM*-mediated regulation of phytohormone pathways, while operating within a drastically reorganized regulatory landscape. Our findings reveal that evolution of the one-leaf developmental system involves rewiring of existing meristematic networks rather than passive suppression of meristem activity.

## Results

### Genome assembly of the allotetraploid *Monophyllaea glabra* confirms the conservation of core meristem regulators

To provide a genomic foundation for investigating the molecular basis of meristem modification in *Monophyllaea*, we generated the first draft genome assembly of *M. glabra* (**Table 1**). The resulting assembly, comprising 181 contigs, is 770 Mb with an N50 of 13.2 Mb. Gene prediction resulted in 70,769 gene models, including 68,490 protein-coding genes. This relatively large number of predicted genes is explained by the tetraploid nature of *M. glabra*, which is further supported by the high proportion of duplicated BUSCOs (63.6%; **Table 1**).

**Table 1.** Summary of *M. glabra* genome assembly.

|  |  |
| --- | --- |
| Total length (bp) | 770,322,622 |
| Number of contigs | 181 |
| Contig N50 (bp) | 13,244,217 |
| Repeat content (%) | 74.2 |
| Predicted genes | 70,769 |
| Protein-coding genes | 68,490 |
| Non-coding genes | 2279 |
| BUSCO complete single copy (%) | 30.7 |
| BUSCO complete duplicated (%) | 63.6 |
| BUSCO fragmented (%) | 2.3 |
| BUSCO missing (%) | 3.4 |

We then performed phylogenetic analysis and divergence time estimation using genomic data from four publicly available Gesneriaceae species (**Figure 1B**). The analyses suggested that divergence between Epithemateae (including the genus *Monophyllaea*) and Trichosporeae (including *Boea* and *Streptocarpus*) tribes occurred 50.0 Mya (45.1–54.4, 95% HPD). We also calculated synonymous substitution rates using four-fold degenerate transversion (4DTv)^31^ for one-to-one ortholog pairs among species and for homeolog pairs within *M. glabra* to estimate the timing of gene duplication (**Figures 1C and 1D**). The peak 4DTv value was 0.018 and the duplication time was estimated to be 3.22 Mya (2.83–3.95, 95% CI). Although further genomic information from the genus *Monophyllaea* is required to evaluate exactly how the genome duplication occurred in the lineage including *M. glabra*, these results suggest that *M. glabra* is an allotetraploid that arose via hybridization between closely related species that diverged ∼3.2 Mya.

The *M. glabra* genome assembly enabled systematic analysis of gene-family evolution and lineage-specific gene losses. Comparative analysis revealed 52 orthologous groups (OGs) that were likely lost in the common ancestor of Trichosporeae. An additional 176 OGs were not detected in any of the currently available Gesneriaceae genomes (**Figure S1A**). Conversely, 975 and 715 OGs were exclusively detected in *M. glabra* and Trichosporeae species, respectively, while 227 OGs were shared between them (**Figure S1B**).

Focusing specifically on *M. glabra*, we found that 326 OGs were undetected, likely representing losses within the Epithemateae tribe, the *Monophyllaea* genus, or *M. glabra* itself (**Figure S1A**). Notably, orthologs typically used as tissue markers in *A. thaliana*, such as *SUCROSE-PROTON SYMPORTER 2* (phloem), *VASCULAR-RELATED NAC-DOMAIN 7* (xylem), and *BLUE LIGHT SIGNALING 1* (guard cell), were undetectable among the gene models. However, inspection of the corresponding gene families indicated that closely related homologs were often retained in *M. glabra* (**Figure S1C**). Thus, the absence of these focal orthologs does not necessarily imply the complete loss of their corresponding physiological functions. Crucially, regarding the morphological divergence of *Monophyllaea*, we investigated the repertoire of orthologs for SAM-related transcription factors. The comprehensive gene annotation revealed that key gene families such as KNOX, CUC, and WUSCHEL-RELATED HOMEOBOX (WOX) are present in the *M. glabra* genome (**Figure 1E**). This finding rules out the possibility that loss of the conventional SAM in the vegetative phase is due to simple gene loss-of-function mutations. This makes complete sense because SAM functions normally during the reproductive phase after floral induction, even in *M. glabra*.

### Tissue-specific transcriptome analysis reveals distinct transcriptional profiles among meristematic regions

To dissect the spatial gene expression dynamics of *M. glabra*, we performed tissue-specific bulk RNA-seq using a microdissection-based tissue isolation method^32^. We isolated five distinct tissues (**Figure 2A**): the GM in the vegetative phase (vGM), the basal and distal regions of the BM (vbBM and vdBM), the microcotyledon (vMIC), and the GM in the reproductive phase (rGM). Using these transcriptomes, we first searched for genes upregulated in the GM relative to other tissues to identify GM-associated genes.

**Figure 2.**
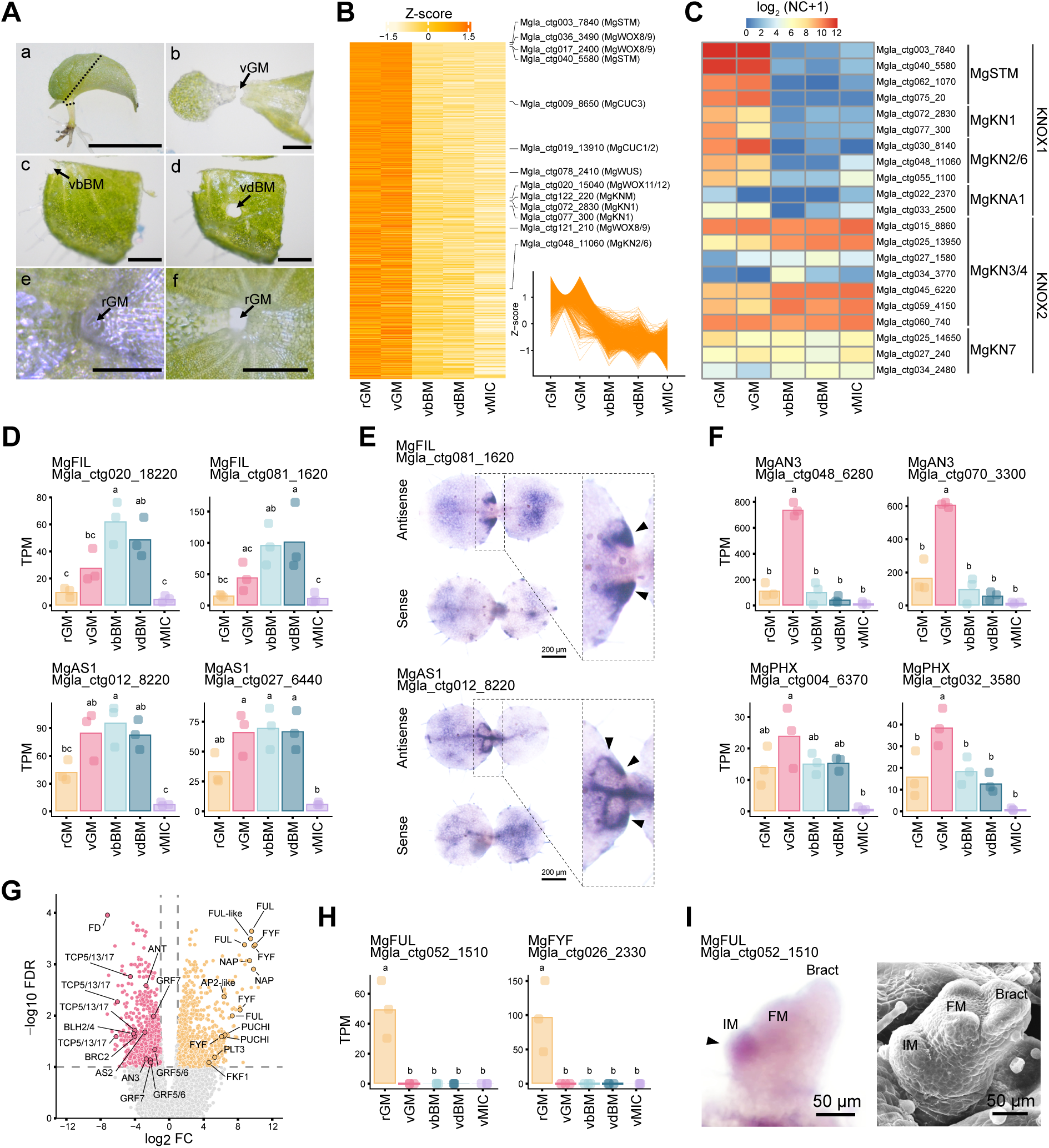
Distinct transcriptional profiles revealed by tissue-specific bulk RNA-seq analysis in *M. glabra*. (A) Microdissection strategy used for tissue-specific RNA-seq. For isolation of vegetative-phase groove meristem (vGM), tissue was excised along the dashed line (a), and the region between the two cotyledons was punched out (b). Basal and distal regions of the basal meristem were separately punched out as vbBM (c) and vdBM (d). Microcotyledon (vMIC) was also punched out. Reproductive-phase groove meristem (rGM), in which a dome-shaped structure is visible (e), was punched out using the same procedure as for vGM (f). Scale bars = 5 mm (a) and 500 µm (b–f). (B) Expression patterns of genes showing higher expression in the rGM and vGM compared with other tissues. Z-scores of normalized counts (NC) are shown. (C) Expression patterns of KNOX1 and KNOX2 genes in *M. glabra*. Values represent log₂-transformed NC + 1. (D, F) Expression patterns of leaf polarity and growth-related genes across tissues. Transcripts per million (TPM) values are shown. Tukey’s HSD test, *p* < 0.001; *n* = 3. (E) Spatial validation of *MgFIL* and *MgAS1* gene expression. Whole-mount *in situ* hybridization (WISH) using antisense (upper) and sense (lower) probes. (G) Differential expression analysis between vGM and rGM. Pink and orange indicate genes upregulated in vGM and rGM, respectively (|log₂FC| > 1, FDR < 0.1; *n* = 3). (H) Representative genes upregulated in rGM. TPM values are shown. Tukey’s HSD test, *p* < 0.001; *n* = 3. (I) WISH showing spatial expression of *MgFUL* in the rGM, alongside a scanning electron microscopy micrograph of the inflorescence apex. IM, inflorescence meristem; FM, floral meristem.

This approach identified several regulators related to shoot formation, including KNOX, CUC, and WOX family genes (**Figure 2B**). Specifically, GM-enriched genes included orthologs of class I KNOX (*MgSTM* and *KNAT1*, hereafter *MgKN1*), *KNAT2* (hereafter *MgKN2*), and class M KNOX (hereafter *MgKNM*).

To determine whether this enrichment reflects functional differences between SAM-maintaining KNOX1 and organ-differentiating KNOX2 genes within the KNOX family^33^, we next performed phylogenetic characterization and expression profiling. These analyses showed that expression of KNOX1 genes was predominantly restricted to the GM and notably absent from the BM, whereas KNOX2 genes exhibited broader expression domains extending into the BM (**Figures 2C and S2**). This pattern strongly indicates that the GM retains core transcriptional features of a conventional SAM.

However, the GM transcriptional landscape was not identical to that of a conventional SAM, consistent with previous observations^3,16^. Leaf lamina program genes such as *FILAMENTOUS FLOWER* and *ASYMMETRIC LEAVES 1* orthologs (*MgFIL* and *MgAS1*) were detected in both the BM and vGM by tissue-specific RNA-seq (**Figure 2D**). To examine these spatial patterns, we performed whole-mount *in situ* hybridization (WISH) analysis. *MgFIL* expression was detected mainly in the BM region, whereas *MgAS1* transcripts were observed in both the GM and BM regions, consistent with the RNA-seq profile for *MgAS1* (**Figure 2E**). Additional leaf morphogenesis regulators such as *MgAN3*, *PHABULOSA/PHAVOLUTA*-related genes (hereafter *MgPHX*), and most *REVOLUTA* orthologs (*MgREV*) were expressed at higher levels in the vGM than in the BM (**Figures 2F and S3**). Together, these results indicate that the vGM retains a SAM-like transcriptional core while expressing multiple components of a leaf lamina program. This overlapping signature was developmentally resolved upon the transition to the reproductive phase, when conventional organogenesis occurs. In the rGM, leaf development and cell proliferation regulators such as the GROWTH-REGULATING FACTOR (GRF) and TCP families were downregulated along with *MgAN3* (**Figure 2G**). Other leaf lamina program genes highly expressed in the vGM (*MgFIL*, *MgAS1*, and *MgPHX*) were also downregulated (**Figures 2D and 2F**). Furthermore, expression of the *FD* ortholog (*MgFD*), a component of the florigen activation complex, was markedly reduced (**Figure 2G**). This pattern is consistent with the downregulation of *FD* in *A. thaliana* floral primordia during the transition from shoot apical to floral meristem identity^34,35^. Concurrently, floral genes such as *FRUITFULL* and *FOREVER YOUNG FLOWER* orthologs (*MgFUL* and *MgFYF*) were upregulated (**Figures 2G–I**). This demonstrates that the GM shifts its transcriptional profile from a chimeric vegetative state to a conventional floral-competent state.

### Single-nucleus RNA-seq reveals cellular heterogeneity within the GM and BM

To resolve the cellular populations underlying the unique morphogenesis of *M. glabra*, we performed droplet-based single-nucleus RNA-seq on the basal region of the macrocotyledon, which encompasses the GM and BM. Samples were collected at 21 days after sowing, when both meristems were fully established (**Figures 1A and 3A**). We profiled 15,313 nuclei from two biological replicates and identified 32 clusters in a uniform manifold approximation and projection (UMAP) space (**Figures 3B and S4; Table S1**). The clusters were consistently represented across replicates, with minimal batch effects (**Figure S4**). We then combined the transcriptomic data with WISH analyses to map selected clusters to their spatial locations within the macrocotyledon (**Figure 3C**). We first examined previously identified GM marker genes to validate the cluster identities^3,16^. *MgSTM* and *MgCUC1/2* were highly enriched in Cluster 23, identifying it as a GM population (**Figure 3D**). *MgAN3* was detected in Cluster 23 and several additional clusters (Clusters 6–8, 11–13, 16, 18, and 19; **Figure 3D**). Its co-detection with *MgSTM* in Cluster 23 extended the tissue-level observation of their overlapping expression to single-nucleus resolution (**Figure 3D**).

**Figure 3.**
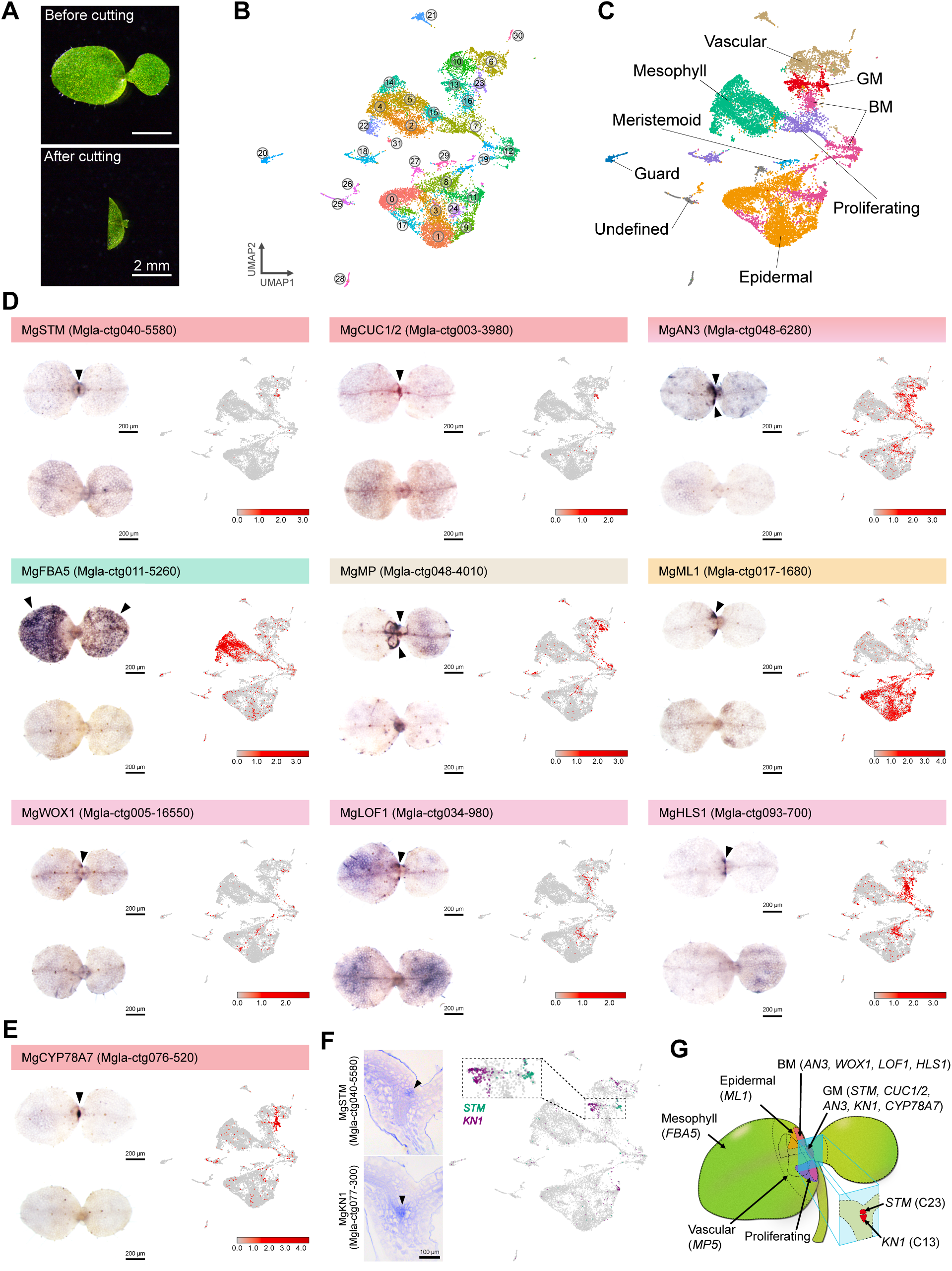
Single-nucleus transcriptomic atlas and spatially informed cell type annotation. (A) Images before and after tissue dissection. The basal region of a 21-day-old seedling was used for snRNA-seq analysis. The basal part of the macrocotyledon encompassing the GM and BM was obtained. Scale bar = 2 mm. (B) Uniform manifold approximation and projection (UMAP) visualization of 15,313 nuclei profiled by snRNA-seq, showing 32 clusters. Each dot represents a single nucleus, colored by cluster identity. (C) UMAP colored by annotated cell types based on marker gene expression and Gene Ontology (GO) enrichment. Distinct colors indicate different cell types. (D) Spatial validation of representative marker gene expression. WISH was performed using antisense (upper left) and sense (lower left) probes, with corresponding gene expression patterns shown on the UMAP (right). (E) Expression pattern of *MgCYP78A7*. WISH results (antisense, upper left; sense, lower left) and UMAP visualization (right). (F) Expression patterns of *MgSTM* and *MgKN1*. WISH staining (left) shows distinct spatial domains within the GM (arrowheads). UMAP visualization (right) highlights nuclei expressing *MgSTM* (green) and *MgKN1* (purple). (G) Schematic summary of tissue organization defining each major cell type identified by snRNA-seq in *M. glabra*.

To place the meristematic clusters within the broader context of the macrocotyledon, we annotated major tissue-specific cell types based on Gene Ontology (GO) enrichment and the orthologs of well-characterized markers (**Figures 3D and S5; Tables S2 and S3**). Orthologs of *FRUCTOSE-BISPHOSPHATE ALDOLASE 5* (*MgFBA5*), *ARABIDOPSIS THALIANA MERISTEM LAYER 1* (hereafter *MgML1*), and

*MONOPTEROS/AUXIN RESPONSE FACTOR 5* (*MgMP*) marked mesophyll, epidermal, and vascular cells, respectively. These markers identified mesophyll Clusters 2, 4, 5, 14, 15, 22, and 31; epidermal Clusters 0, 1, 3, 8, 9, 11, 17, 24, 26, and 29; and vascular Clusters 6 and 21 (**Figure 3D**). Additional clusters were annotated as proliferating (Clusters 7 and 18), residual vascular (Clusters 6, 10, 21, and 30), meristemoid (Cluster 29), or guard cell populations (Cluster 20) based on GO enrichment (**Tables S2 and S3; Figure S5**).

We next characterized BM-associated populations using orthologs of *WOX1* (*MgWOX1*) as a leaf margin marker and *LATERAL ORGAN FUSION 1* (*MgLOF1*) as a boundary region marker (**Figure 3D**). These markers identified Clusters 7, 8, 11, 12, 16, 17, and 19 as candidate BM clusters. *HOOKLESS 1* ortholog (*MgHLS1*), which accumulated in the BM and was highly expressed in Clusters 11 and 16, further supported these assignments (**Figure 3D**). We therefore annotated Clusters 11, 12, 16, 17, and 19 as BM cells, whereas Clusters 7 and 8 were classified as proliferating and epidermal populations, respectively.

Finally, we investigated the GM to explore its spatial and transcriptional complexity. Alongside the previously identified markers, expression of the *CYP78A7* ortholog (*MgCYP78A7*) further supported the assignment of Cluster 23 as a GM population (**Figure 3E; Table S3**). Unexpectedly, while our tissue-specific transcriptome analysis identified *MgKN1* as a GM-enriched gene (**Figure 2C**), it was not co-expressed with *MgSTM* in Cluster 23 (**Figure 3F; Table S3**). Instead, *MgKN1*-expressing cells were highly enriched in Cluster 13. Spatial analysis further showed that *MgKN1* expression was restricted to the hypocotyl-facing side of the GM and did not overlap with the *MgSTM*-expressing domain (**Figure 3F**). This finding demonstrated that the GM was spatially subdivided into two distinct transcriptional domains, which could not be resolved by tissue-level analysis. Taken together, these complementary insights revealed the spatial architecture and cellular diversity of the GM and BM, as summarized in the schematic model in **Figure 3G**.

### Distinct phytohormone regulatory programs in GM and BM

After annotating GM-associated populations, we examined their potential functions through GO enrichment analysis (**Figure S5B**). Notably, terms related to CK dehydrogenase activity were significantly enriched in the GM clusters, suggesting active regulation of CK homeostasis rather than simple maintenance of high CK levels.

We therefore analyzed the expression of phytohormone biosynthesis and inactivation genes across annotated populations (**Figure 4A**). In the BM, orthologs of genes involved in auxin and brassinosteroid biosynthesis, including YUCCA (YUC), CONSTITUTIVE PHOTOMORPHOGENIC DWARF (CPD), and DWARF 4 (DWF4) family members, were expressed (**Figures 4A and S6**). In contrast, GM cell populations expressed orthologs of genes involved in CK (e.g., *MgCYP735*, *MgIPT*, and *MgLOG*) and brassinosteroid (e.g., *MgBR6ox*) biosynthesis (**Figures 4A and S6**), together with genes involved in the inactivation of CK (*Cytokinin oxidase/dehydrogenase*: *MgCKX*), GA (*MgGA2ox*), and auxin (*MgGH3*). In particular, multiple CKX family genes, such as *CKX1* orthologs (*MgCKX1*), were detected in the GM (**Figure S7**). These expression patterns indicate that GM and BM operate distinct phytohormone regulatory programs.

**Figure 4.**
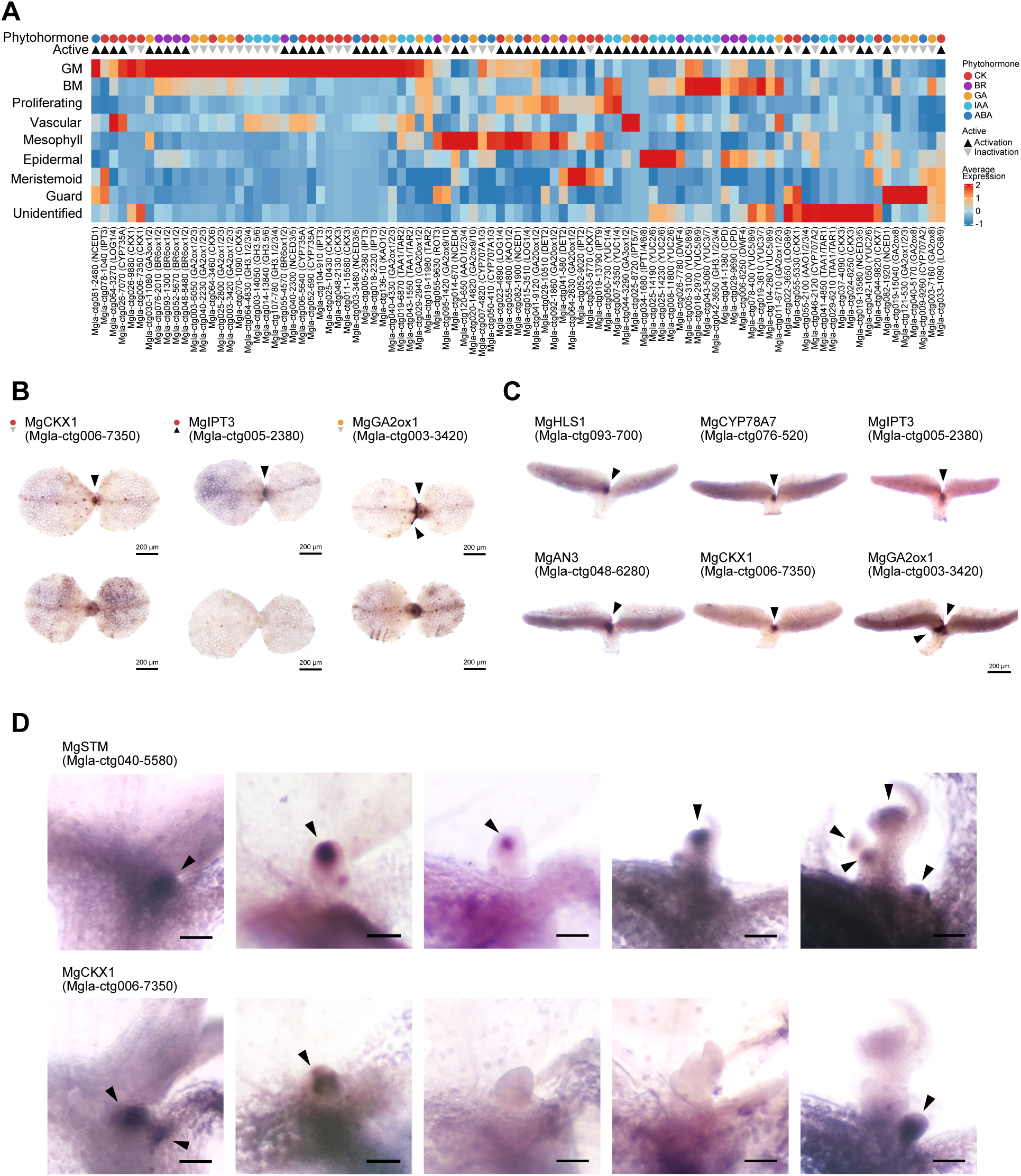
Differential phytohormone regulation between the GM and BM. (A) Pseudo-bulk average expression of phytohormone-related genes across annotated cell types derived from snRNA-seq. Genes involved in phytohormone biosynthesis (black triangles) and inactivation (gray inverted triangles) are shown for each phytohormone class. Colors indicate phytohormone categories: cytokinin (CK, red), brassinosteroid (BR, purple), gibberellin (GA, orange), auxin (IAA, light blue), and abscisic acid (ABA, blue). (B) Spatial validation of representative phytohormone metabolic genes by WISH. Expression patterns of the CK inactivation gene *MgCKX1*, the CK biosynthesis gene *MgIPT3*, and the GA inactivation gene *MgGA2ox1* are shown using antisense (upper) and sense (lower) probes. Scale bars = 200 µm. (C) Side views of WISH samples highlighting spatial biases in gene expression within the GM and surrounding tissues. Arrowheads indicate gene expression domains revealed by staining. Scale bars = 200 µm. (D) Spatial expression patterns of *MgSTM* and *MgCKX1* revealed by WISH. Panels from left to right show sequential developmental stages: the transition phase from the vegetative GM to inflorescence meristem (IM) formation, the establishment of a dome-shaped IM, the initiation of bract formation, the initiation of the next IM, and the clear visualization of the subsequent IM. Scale bars = 100 µm.

We next validated these phytohormone-metabolic features by examining spatial expression patterns using WISH (**Figure 4B**). Both the CK biosynthesis gene *MgIPT3* and the CK inactivation gene *MgCKX1* were expressed in the GM, while the ortholog of the GA-inactivating gene *MgGA2ox1* was detected in the GM as well as along the basal margin of the macrocotyledon (**Figure 4B**). In contrast to the canonical SAM, where high CK and low GA levels are maintained^21–23^, the GM of *M. glabra* appears to integrate an additional CK inactivation pathway with the conserved biosynthesis machinery, in parallel with GA inactivation.

A detailed examination of these spatial profiles revealed that the transcripts, while largely expressed within the GM, exhibited spatially biased localization patterns (**Figure 4C**). Specifically, while *MgHLS1* expression was restricted to the BM, *MgAN3* spanned both the BM and the macrocotyledon-facing side of the GM. Within the GM, *MgCYP78A7* expression overlapped with *MgAN3* but was shifted closer to the microcotyledon. The phytohormone metabolic genes also exhibited spatial biases: within the GM, the expression domains of *MgCKX1* and *MgIPT3* overlapped with that of *MgCYP78A7* but were shifted toward the microcotyledon, whereas *MgGA2ox1* was expressed in the GM and extended into the GM-hypocotyl junctions (**Figure 4C**). These partially biased expression patterns suggest that the two GM-associated cell populations resolved by snRNA-seq could be further subdivided into distinct hormone regulatory subdomains.

To further investigate the role of CK regulation, we monitored *MgCKX1* expression during the transition from the vegetative to the reproductive phase (**Figure 4D**). While *MgSTM* marked the center of the dome-shaped inflorescence and floral meristems, *MgCKX1* exhibited a distinct, compartmentalized expression pattern and was specifically localized to the region between the inflorescence meristem and the microcotyledon. Unlike the stable expression of *MgSTM*, *MgCKX1* expression was transient: it became undetectable as the structure developed but reappeared in the primordia of the emerging inflorescence axis (**Figure 4D**). This mutually exclusive and transient spatial arrangement suggests that localized CK inactivation by MgCKX1 creates a low-CK zone that partitions the active meristem from surrounding tissues during dome formation.

### GM cell populations exhibit SAM-like identities with unique regulatory programs

To characterize the distinctive features of the GM, we next compared it with the canonical dome-shaped SAM of *A. thaliana* (**Figure 5**). We integrated our *M. glabra* snRNA-seq dataset with reannotated *A. thaliana* single-cell RNA-seq (scRNA-seq) data^36^ (**Figures 5A and 5B**). As an internal positive control for the integration, we confirmed that major tissue-specific populations aligned across species: *M. glabra* mesophyll, vascular, meristemoid, and guard cells clustered with their *A. thaliana* counterparts (**Figures 5C and S8**). We also noted that certain cell populations remained unaligned, confirming the preservation of species- or developmental stage-specific identities (**Figure S8**). Within this integrated space, BM-associated cells were distributed across clusters enriched for proliferating and epidermal cells, consistent with their indeterminate nature (**Figure 5C**).

**Figure 5.**
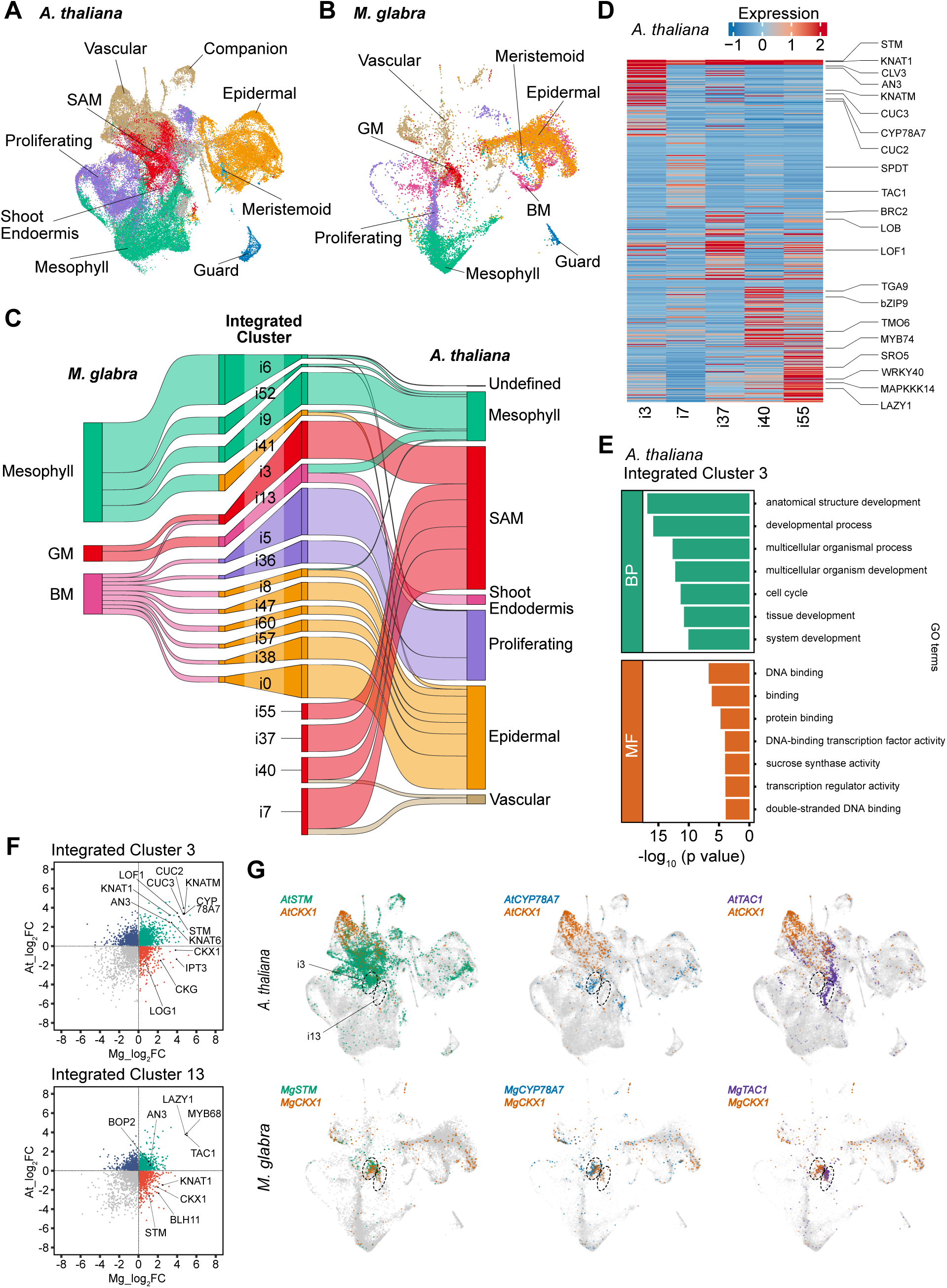
GM-specific traits highlighted by cross-species integration with *A. thaliana*. (A, B) UMAP visualization of clusters from the *A. thaliana* shoot apex (A) and the basal region of *M. glabra* macrocotyledons (B) following cross-species integration analysis. Cells are colored by integrated cluster identity. (C) Sankey diagram illustrating correspondence between annotated cell types of *M. glabra* and *A. thaliana* based on integrated clusters. A comprehensive view is shown in **Figure S8**. (D) Pseudo-bulk average expression profiles of representative genes across *A. thaliana* shoot apical meristem (SAM)-related integrated clusters, highlighting transcriptional differences among SAM subpopulations. (E) GO enrichment analysis of genes enriched in the *A. thaliana* SAM-associated integrated cluster i3. Biological Process (BP) and Molecular Function (MF) terms are shown. (F) Cross-species comparison of gene enrichment in integrated clusters i3 and i13. The x-axis (Mg_log_2_FC) indicates log_2_ fold changes of genes in i3 (or i13) relative to other *M. glabra* clusters, and the y-axis (At_log_2_FC) indicates log_2_ fold changes relative to other *A. thaliana* clusters. Green dots represent genes enriched in both species, red dots represent genes enriched in *M. glabra* but depleted in *A. thaliana*, and blue dots represent genes depleted in *M. glabra* but enriched in *A. thaliana*. (G) Integrated UMAP visualization highlighting nuclei expressing *STM* (green), *CYP78A7* (blue), and *TAC1* (purple) in *A. thaliana* and *M. glabra*.

In contrast, GM-associated cells mapped primarily to two specific integrated clusters, i3 and i13 (**Figure 5C**). Based on *A. thaliana* annotations, i3 corresponds to SAM cells, while i13 represents shoot endodermis cells implicated in gravitropism. Accordingly, canonical meristem genes such as *STM* and *KNAT1* (*MgKN1*) were primarily expressed in i3, with minimal detection in i13. Notably, whereas *A. thaliana* SAM cells were dispersed across multiple clusters (i3, i7, i37, i40, and i55), *M. glabra* GM cells were selectively concentrated in i3 among those representing *A. thaliana* SAM cells (**Figure 5C**). Differential expression analysis in *A. thaliana* revealed that i3 represents the morphogenetically active SAM core, distinguished by the enrichment of meristematic maintenance and boundary genes (*AtCLV3*, *AtCUC*) rather than by the vascular or gravitropic signatures (*AtTAC1*, *AtLAZY1*) found in other *A. thaliana* SAM-associated clusters (**Figures 5D and 5E**). Although the cross-species integration lacked the fine resolution to separate the *MgSTM*- and *MgKN1*-expressing GM subpopulations observed in our single-species analysis (**Figure 3F**), this major GM population specifically aligned with the cell population governing meristem maintenance within the SAM.

Building on this conserved core identity, we investigated the molecular basis of the morphological divergence by comparing gene expression within these clusters (**Figure 5F**). In i3, while *STM* and *KNAT1* (*MgKN1*) were shared between species as described above, *M. glabra* cells displayed a distinct enrichment of CK-related genes, including *CKX1* and *IPT3* (**Figure 5F**). Similarly, in the shoot endodermis integrated cluster i13, *CKX1* and *STM* were highly expressed in *M. glabra* despite the conservation of gravitropic markers (*TAC1*, *LAZY1*; **Figure 5F**). Crucially, this difference extended to cellular distribution. While *STM* and *CKX1* were largely expressed in mutually exclusive cells in *A. thaliana*, their expression overlapped within integrated cluster i3 in *M. glabra* (**Figure 5G**). These results indicate that the GM retains a core SAM identity but uniquely integrates negative regulators of CKs into the same cellular context, establishing an intracellular chimeric identity that likely underlies its distinct morphology.

## Discussion

Our study revealed that the unique one-leaf body plan of *M. glabra* arises through reorganization of the meristematic system rather than loss of canonical SAM components or passive arrest of SAM activity. Morphologically, the *M. glabra* GM has long been hypothesized to resemble the *A. thaliana stm* mutant, based on its loss of the dome-shaped morphology and limited organogenic activity^15,17^. However, our genomic analysis confirmed the retention of core SAM regulators, including the STM, CUC, and WUS gene families (**Figure 1E**), indicating that the unusual GM morphology cannot be explained simply by the absence of these core regulator families. Although the same analysis identified *M. glabra* as an allotetraploid, the associated hybridization event itself is unlikely to be the primary developmental trigger. Indeed, homologous gene pairs encoding key regulators generally exhibited similar expression patterns without obvious subgenome dominance (e.g., **Figures 2D and 2F**), supporting the idea that the unique morphology is attributable to transcriptional rewiring rather than genomic asymmetry.

A striking feature of this rewiring is altered phytohormonal regulation (**Figure 4**). In the *A. thaliana* SAM, CK signaling promotes *WUS* expression to maintain the stem cell niche^25^. Downregulation of active CK accumulation by *CKX1* overexpression reduces SAM size^37–39^, whereas overaccumulation of active CK in *ckx3 ckx5* mutants results in an enlarged SAM^40^. Thus, CKX-mediated CK status determines the size and activity of the meristematic center. Consistent with this role, several *CKX* genes are expressed in the shoot apex of *A. thaliana*, though their expression profiles can vary across developmental stages or be spatially restricted around the SAM^39–41^. *CKX3* is expressed in the center of the inflorescence meristem, but it is not expressed in certain SAM stages^40^. Similarly, *CKX5* is expressed around the SAM but is strongly expressed in the procambium rather than the meristematic center^40^. Furthermore, while *CKX1* expression in the shoot apex was previously detected by promoter-β-glucuronidase analysis^39^, our reanalysis of *A. thaliana* scRNA-seq data from the vegetative shoot apex^36^ revealed that *CKX1* and *STM* were expressed in distinct cell populations, suggesting that *CKX1* is predominantly localized outside the central niche (**Figure 5**). Consequently, unlike in *A. thaliana*, where *CKX* expression is spatially excluded from the stem cell niche or temporally regulated, the GM integrates *CKX*-mediated inactivation into its core program (**Figures 4 and 5**). The concurrent expression of *MgIPT3* and *MgCKX1* in GM cells likely establishes a unique homeostatic balance that acts as a constitutive brake, mimicking the *A. thaliana CKX1* overexpression phenotype (**Figure 4**)^37,38^. This balance likely underlies the persistently low organogenic activity of the vegetative GM^3^, preventing dome formation while retaining the capacity for future reproductive development.

The GM also exhibits a transcriptional identity distinct from both the conventional SAM and the BM. The BM is characterized by the distinct expression of phytohormonal biosynthesis genes and leaf lamina program genes such as *MgAS1*, *MgFIL*, and *MgWOX1*, whereas KNOX1 expression is restricted to the GM (**Figures 2–4 and S2**). Nevertheless, the GM also expresses leaf-associated regulators including *MgAN3* and *MgAS1*, indicating that a leaf lamina program is superimposed on a conserved SAM network. This chimeric transcriptional state brings together regulatory components that are normally separated or antagonistic. KNOX1 genes promote CK biosynthesis^21,22^. The GRF-GIF/AN3 module enhances CK signaling in *A. thaliana* and represses *CKX* expression in poplar^42,43^. In contrast, *AS1* represses KNOX1 genes such as *STM*, potentially limiting the CK-promoting activity^44,45^. The concurrent expression of *MgCUC* further amplifies this chimeric state (**Figure 3**)^16^. The overlap of *MgSTM* and *MgCUC* expression is reminiscent of an early embryonic stage, prior to the separation of the SAM from cotyledonary domains^30^. However, the simultaneous presence of post-embryonic leaf-associated regulators suggests that the GM is not merely a passive arrest of an embryonic state. Instead, *M. glabra* appears to actively maintain an immature, plastic state by integrating multiple, often conflicting, developmental modules.

This chimeric transcriptional state changes during the transition to reproductive development. Leaf lamina program genes that are highly expressed in the vegetative GM are downregulated in the rGM, whereas floral regulators are upregulated (**Figure 2G**). These temporal changes indicate that incorporation of a leaf lamina program is primarily associated with the vegetative groove state. However, *MgCKX1* expression persists during dome formation and becomes localized to boundary regions of the reproductive meristem (**Figure 4D**), suggesting that MgCKX1 performs an additional role in compartmentalizing the emerging inflorescence. Thus, the one-leaf phenotype likely results from a specific combination of SAM identity, leaf-associated transcriptional activity, and phytohormone regulation, rather than from any single regulatory alteration. Further single-cell analysis across the vegetative-to-reproductive transition will be required to distinguish the mechanisms that maintain the groove state from those that promote dome emergence, although this is beyond the scope of this study.

Cross-species analysis at single-cell resolution clarifies the evolutionary implications of the GM. Despite the evolutionary divergence between *A. thaliana* and *M. glabra*, GM cells aligned with the conserved SAM core represented by integrated cluster i3 (**Figure 5C**). Our conservative integration retained several species-specific, unaligned populations, and the alignment therefore indicates similarity in broad cell-state identity rather than transcriptome-wide equivalence (**Figure S8**). Within the aligned SAM population, specific regulatory components, particularly CK-related genes such as *CKX1*, differed between the two species (**Figures 5F and 5G**). Although further validation is required, the coexistence of a conserved SAM-like cellular identity and species-specific regulatory programs supports the interpretation that GM cells evolved through the co-option and reorganization of pre-existing SAM-, leaf-, and embryo-associated developmental modules rather than as a completely novel cell type.

The *de novo M. glabra* genome assembly, the first reported for the tribe Epithemateae, also provides a phylogenetic reference for comparative studies across Gesneriaceae. Because Epithemateae includes genera with conventional shoot architectures (e.g., *Epithema*, *Rhynchoglossum*, and *Whytockia*)^10^, comparative analysis using the *M. glabra* genome and additional Epithemateae genomes will enable the identification of genomic changes specifically associated with the unique morphology of *Monophyllaea*. As an outgroup to the well-sequenced sister tribe Trichosporeae^46–49^, the *M. glabra* genome also helps resolve ancestral genomic states and identify evolutionary events that occurred in the common ancestor of Trichosporeae (**Figure S1**). Crucially, extreme one-leaf morphologies have also evolved independently within Trichosporeae (e.g., in certain *Streptocarpus* species)^27^. The genomic baseline established here thus sets the stage for future comparative studies across Epithemateae to elucidate the genomic basis of the parallel evolution of this striking body plan across distinct lineages.

In summary, our combined spatial and single-nucleus analyses provide a molecular framework for the fuzzy morphology of *Monophyllaea*. Partial morphological overlap between leaf and shoot is accompanied by the coexistence of SAM-associated, leaf-associated, embryo-associated, and phytohormone regulatory programs within the GM. Their reorganization at the cellular level illustrates how conserved developmental modules can generate novel morphologies and contribute to plant diversity.

## Resource availability Lead contact

Further information and requests for resources and reagents should be directed to and will be fulfilled by the lead contact, HT.

## Materials availability

This study did not generate new, unique reagents.

## Data and code availability

`o`Raw RNA-seq data have been deposited in the DNA Data Bank of Japan (DDBJ) Sequence Read Archive (DRA) under BioProject accession number: PRJDB42067.
`o`The complete analytical pipeline for snRNA-seq analysis will be made publicly available upon publication (https://github.com/Shunji-Nakamura/Research-paper-for-snRNAseq-of-Monophyllaea).
`o`Any additional information required to reanalyze the data reported in this paper is available from the lead contact upon request.

## Acknowledgments

We thank Drs. Keiji Nakajima and Tatsuaki Goh for experimental support and insightful discussions regarding plant nuclei isolation, Dr. Junko Watanabe for assistance with nuclei sorting, and Drs. Akira Nagatani, Nobuyoshi Mochizuki, and Sujung Kim for assistance with tissue isolation. This work was supported by JSPS KAKENHI Grant Numbers JP25KJ0072 (to S.N.), JP19J14140 (to A.K.), JP25113002 (to H.T.), JP19H05672 (to H.T.), and JP24H00566 (to H.T.). S.N. was additionally supported by a Sasakawa Scientific Research Grant from The Japan Science Society (2023−5017).

## Author contributions

S.N. and H.T. conceived and designed the research. A.K. conducted plant cultivation and tissue sampling for genome sequencing and bulk RNA-seq. H.K. established the draft genome assembly and led the evolutionary genomic analyses. S.N. and H.K. processed and analyzed the bulk RNA-seq data. S.N. generated the snRNA-seq atlas (including nuclei preparation, sorting, and bioinformatics) and performed the spatial gene expression analyses (WISH) alongside developmental profiling. S.N. and H.T. wrote the manuscript. All authors discussed the results and edited and approved the final manuscript.

## Materials and Methods

### Plant materials and growth conditions

*M. glabra* seeds were originally collected at Srakaew Cave, Thailand in January 2000, with permission from Mr. T. Wongprasert of the Forest Herbarium, Thailand. Voucher specimens are housed in the Forest Herbarium, Department of National Parks, Wildlife and Plant Conservation, Bangkok (BKF), and the University of Tokyo Herbarium (TI). The strain was maintained by cultivation in growth chambers. Plants were grown in plates containing one-third-strength Murashige and Skoog (MS) salts and 0.8% agar at 22°C under short-day (SD; 8 h light and 16 h dark) conditions. For tissue-specific bulk RNA-seq experiments during the vegetative phase, plants were also grown under continuous light (CL) conditions with a light intensity of ∼45 µmol m^-2^ s^-1^.

### Genome sequencing

*M. glabra* plants grown for 16 days on one-third-strength MS medium were placed in 2.0 mL tubes containing 5 mm and 0.5 mm zirconia beads and immediately frozen in liquid nitrogen. Plants were homogenized by shaking with a TissueLyser II (Qiagen, Hilden, Germany) at 30 rps for 3 min. Cetyltrimethylammonium bromide (CTAB) buffer (1.0 mL; 2% w/v CTAB, 2% w/v PVP-30, 100 mM Tris-HCl pH 8.0, 25 mM ethylenediaminetetraacetic acid, 2 M NaCl) was added to the homogenate, incubated at 50°C for 30 min, and centrifuged at 8000 × *g* for 10 min. The supernatant was transferred to a new tube, and RNase A (0.35 mg) was added and incubated at 37°C for 30 min. Chloroform (1.0 mL) was added and mixed thoroughly, centrifuged at 8000 × g for 5 min, and the aqueous phase was transferred to a new tube. This extraction was repeated three times, but in the second extraction phenol:chloroform:isoamyl alcohol (PCI) was used instead of chloroform. DNA was precipitated by adding an equal volume of 2-propanol and spooled with a glass rod. After washing with 70% (v/v) ethanol, DNA was dissolved in elution buffer (10 mM Tris-HCl pH 7.6).

Long-read sequencing was performed on a MinION platform from Oxford Nanopore Technologies (ONT; Oxford, UK). Libraries were prepared using a ligation-based kit (SQK-LSK110) and sequenced on two flow cells (R9.4.1). Base calling was performed with Guppy (v5.0.15; ONT) in super accurate mode, yielding 30.1 Gb of long reads. Short-read libraries were prepared using a Collibri PCR-free ES DNA Library Prep Kit for Illumina Systems (Thermo Fisher Scientific, Waltham, Massachusetts, USA) and sequenced by Macrogen Japan using a HiSeq X Ten instrument to generate 150 bp paired-end reads. This yielded 57.6 Gb of short reads. Sequence reads have been deposited in the DNA Data Bank of Japan (DDBJ) Sequence Read Archive (DRA).

### Genome assembly

The *M. glabra* predicted genome size was 637.8 to 844.6 Mb according to k-mer-based estimation. ONT long reads were adapter-trimmed using Porechop v0.2.4 (https://github.com/rrwick/porechop) and quality/length filtered with NanoFilt v2.8.0^50^ (q ≥ 7, l ≥ 1000 bp). Illumina short reads were quality-controlled using fastp v0.22.0^51^. *De novo* assembly was performed with NECAT v0.0.1^52^ using the filtered ONT reads. The draft assembly was polished with ONT reads by two rounds of Racon v1.4.3^53^, followed by Medaka v1.3.2 (model r941_min_high_g360; https://github.com/nanoporetech/medaka), then further polished with Illumina reads using hypo v1.0.3^54^, where short reads were mapped to the assembly using minimap2 v2.22^55^ with secondary alignments disabled. Redundant haplotigs were evaluated using purge_haplotigs^56^, and purging was performed with “-a 99” to preferentially remove contigs predicted as artifacts while retaining primary contigs. The final polished assembly consisted of 181 contigs with an N50 of 13.2 Mb (**Table 1**).

### Bulk RNA-seq for gene prediction

For gene prediction, vegetative seedlings at 20 days after sowing were collected along with inflorescences from plants. Total RNA was extracted using a Plant Mini Kit (Qiagen) with DNase I treatment. RNA-seq libraries were prepared using a KAPA Stranded mRNA-seq Kit (KAPA Biosystems, Wilmington, MA, USA) according to the manufacturer’s protocol. Sequencing was performed on an Illumina HiSeq 1500 platform (Illumina) to generate 150 bp paired-end reads.

### Gene prediction

RNA-seq reads were quality-controlled using fastp v0.23.2 and aligned to the genome assembly using STAR v2.7.9^57^. Splice junctions supported by the RNA-seq alignments were curated using Portcullis v1.2.4^58^, which analyzes splice junctions in BAM files and filters junctions that are unlikely to be genuine. Repetitive elements were identified with RepeatModeler v2.0.3^59^ and masked using RepeatMasker v4.1.2-p1 (http://www.repeatmasker.org) prior to gene prediction.

Gene models were then generated using three complementary approaches and integrated. First, BRAKER2 v2.1.6^60–63^ was used for *ab initio* prediction on the repeat-masked genome, running three configurations that incorporated (i) RNA-seq-derived hints, (ii) protein hints from OrthoDB^64^ Plants, and (iii) hints derived from the RNA-seq-based transcript models described below. Second, homology-based prediction was performed with GeMoMa v1.8^65,66^, using the RNA-seq BAM file together with protein datasets from *A. thaliana*, *Solanum lycopersicum*, *Mimulus guttatus*, and *Streptocarpus rexii* (**Table S4**). Third, an RNA-seq-driven transcriptome annotation was constructed by aligning transcript assemblies from StringTie2 v2.2.1^67^ and genome-guided Trinity v2.14.0^68^ to the genome using PASA v2.5.2^69^, and combining these with independently generated genome-guided transcripts from Scallop v0.10.5^70^ and Strawberry v1.1.2^71^. These transcript evidence sources were integrated by Mikado v2.3.3^72^ to select an optimized, non-redundant set of RNA-seq-based models.

Predictions from the three approaches were combined with EvidenceModeler (EVM) v1.1.1^73^ to produce a unified set of coding gene models. To resolve discrepancies between the EVM models and the Mikado transcript set, they were compared using gffcompare v0.11.2^74^; in cases showing major inconsistency, the RNA-seq-based Mikado model was preferentially adopted. Gene models overlapping annotated repeats by ≥ 50% were removed. Finally, PASA annotation updates were performed for three iterative rounds to refine gene structures and add untranslated regions (UTRs), and Mikado-predicted non-coding transcript models were incorporated into the final annotation set. Proteome completeness of the predicted gene set was assessed using BUSCO v6.0.0^75^ with the eudicotyledons_odb12 dataset.

### Orthology assignment

Orthology inference was performed independently using two complementary species sets. For detailed gene family evolution analyses, we compiled a dataset including all publicly available genomes from Gesneriaceae species (**Table S4**) and inferred orthogroups and orthologous relationships using OrthoFinder v3.0.0^76^ with default settings unless otherwise noted. In parallel, to support downstream expression analyses that require robust orthology links to well-annotated reference species, we constructed a model-species-centered dataset and ran OrthoFinder v2.5.5^77^ to infer orthology relationships between *M. glabra* and representative model plants.

### Divergence time estimation

Due to a lack of chromosomal phasing information for *M. glabra*, we could not assign contigs to each subgenome. Accordingly, we determined the WGD-derived duplicated gene pairs of coding genes based on gene collinearity and orthology. For phylogenetic estimation, we constructed a pseudo-diploid proteome for tetraploid species (*M. glabra*, *Oreocharis mileensis*, and *S. rexii*) by randomly retaining one gene of a WGD-derived duplicate. Using the pseudo-diploid proteome data, we performed OrthoFinder analysis and obtained 1727 single-copy orthologous groups that were represented in at least seven species.

Divergence-time estimation was performed with MCMCtree in PAML (v4.10.9)^78,79^. Protein sequences of orthogroups were aligned with MAFFT v7.525^80^, trimmed with ClipKIT (smart-gap, v2.7.0)^81^, and concatenated into a supermatrix. Gene trees were inferred with IQ-TREE v3.0.1^82^ under LG+R4, and a species tree was estimated with ASTRAL v1.23.3.7^83^; this tree was rooted using *A. thaliana* as the outgroup and used as the MCMCtree topology. Time calibrations were set from TimeTree 5^84^ for the *A. thaliana*-*S. lycopersicum* split (111.4–123.9 Ma) and the *S. lycopersicum*-*A. majus* split (75.8–96.6), with an additional soft upper constraint on root age. For molecular dating, we used the amino acid LG model with the independent rates relaxed clock (clock = 2). Following the approximate likelihood workflow, branch-length gradients/Hessian were generated and used in the final dating run.

*Ks* and 4DTv analyses were performed for interspecific comparisons and for within-genome *M. glabra* duplicated pairs. Interspecific gene pairs were derived from OrthoFinder orthologue tables by retaining only strict one-to-one pairs for interspecific comparison. Protein pairs were aligned with MAFFT v7.525, and codon alignments were generated with pal2nal v14.1^85^. 4DTv was calculated from codon alignments using an in-house script. Pairwise 4DTv data were filtered by alignment quality, retaining pairs with no stop/ambiguous sites, alignment length ≥ 100, and gap-site fraction ≤ 0.5. For each dataset, 4DTv distributions were fitted with Gaussian mixture models, and one representative peak was selected using an alignment quality score. Calibration used MCMCtree posterior node ages. Selected interspecific peaks were mapped to their corresponding divergence ages, and a through-origin model was fitted. The *M. glabra* duplicated pair peak was then projected onto this calibration to estimate divergence time. Uncertainty was propagated by bootstrap resampling of peak positions and Monte Carlo sampling of MCMCtree posterior ages, and the final estimate was summarized as median and 95% CI.

### Tissue-specific bulk RNA-seq: sampling, library preparation, and data processing

To characterize tissue-specific expression patterns, vegetative individuals grown under CL conditions for 37–39 days were collected along with reproductive individuals grown under SD conditions for 38 days. Five distinct tissues were excised by punching out with a microdissection instrument^32^: the groove meristem (vGM), the basal part of the basal meristem (vbBM), the distal part of the basal meristem (vdBM), and microcotyledons (vMIC) from vegetative individuals, as well as the emerging dome-shaped groove meristem (rGM) from reproductive individuals. Each biological replicate comprised pooled tissues from 10 individuals, and three biological replicates were prepared per tissue type. The isolated tissues were collected into tubes containing zirconium beads and 50 µL of RLT buffer (Qiagen) supplemented with 2% v/v 2-mercaptoethanol, and immediately frozen in liquid nitrogen. The tissues were homogenized by vigorous shaking using a Mixer Mill M300 (Retsch) at 25 Hz for 1 min, and this procedure was repeated once. Reverse transcription and whole-transcript amplification (WTA) were performed according to the “Whole-transcript amplification for population Quartz-Seq (vPurifiedRNA)” protocol^86^, with 15 cycles of PCR conducted for the WTA step. Sequencing libraries were prepared using a KAPA HyperPlus Kit (KAPA Biosystems) following the manufacturer’s protocol, which was optimized to produce ∼200 bp inserts. Sequencing was performed on an Illumina HiSeq 1500 platform (Illumina) in single-read, rapid-run mode.

All raw sequencing reads were processed using fastp v0.23.2 to remove the reverse transcription, tagging, and WTA primers associated with the Quartz-Seq protocol (**Table S5**). Trimmed reads were mapped to the reference genome using STAR v2.7.10b, and transcript abundances were subsequently quantified using RSEM v1.3.1^87^.

### Identification of tissue-specific expression patterns

Using the raw read counts generated for the five tissue types, genes with low expression levels were filtered out using the filterLowCountGenes function (threshold count = 3) in the R package TCC v1.46.0^88^. To obtain robust normalization factors, the multi-step normalization method implemented in TCC was applied based on the Trimmed Mean of M-values (TMM) method within the edgeR v4.4.0^89^ framework. Iterative normalization was performed using three iterations, a false discovery rate (FDR) of 0.1, and a floorPDEG (proportion of differentially expressed genes) of 0.05.

To identify genes with tissue-specific expression profiles, particularly those specific to the groove meristem (GM), we utilized the empirical Bayesian approach implemented in the R package baySeq v2.28.0^90^. The normalized library sizes derived from TCC were incorporated into the baySeq countData object. We defined four empirical models of expression patterns across the samples: (1) NDE (non-differentially expressed across all samples), (2) GM-specific (vGM and rGM vs. others), (3) BM-specific (vbBM and vdBM vs. others), and (4) All (distinct expression in each tissue type). Prior distributions were estimated using a negative binomial (NB) method with quasi-likelihood (QL) estimation and a sample size of 100,000. Subsequently, posterior likelihoods for each model were calculated using the Bayesian Information Criterion (BIC). Genes were assigned to the expression pattern that yielded the highest posterior probability. For visualization of GM-specific gene expression, normalized counts were log2-transformed or converted to Z-scores, and heatmaps were generated using the R package ComplexHeatmap^91^.

### Differential expression analysis between vegetative and reproductive GMs

To characterize transcriptomic changes during the developmental transition of the meristem, a pairwise differential gene expression analysis was conducted between vGM and rGM samples. Normalization and DEG identification were performed using the TCC package with the edgeR v4.0.16 test method. Genes were considered significantly differentially expressed if they met the criteria of FDR (*q*-value) < 0.05 and absolute log2 fold change (|log2FC|) ≥ 1. The results were visualized as a volcano plot using the ggplot2 and ggrepel packages, highlighting significantly upregulated and downregulated genes along with specific markers of interest.

### Molecular phylogenetic analyses

We retrieved sequences with high similarity using BLASTp against the customized database in which the *M. glabra* proteome and public proteome datasets collected from Phytozome (https://phytozome-next.jgi.doe.gov/) and other sources were integrated (**Table S4**). Sequences were aligned with MAFFT v7.480, and possible non-homologous sites were trimmed from the alignment by TrimAL v1.4^92^. Maximum likelihood trees were reconstructed by IQ-TREE v2.31.1^93^.

### Whole-mount *in situ* hybridization

cDNA sequences were amplified using primers for target genes (**Table S6**). Amplified fragments were cloned into the pZErO-2 vector. Templates for probe synthesis were amplified by PCR using M13 forward (5′-GTA AAA CGA CGG CCAGT-3′) and M13 reverse (5′-CAG GAA ACA GCT ATGAC-3′) primers. Using the templates, we prepared digoxigenin (DIG)-labeled antisense and sense probes using SP6 or T7 polymerases (Roche, Basel, Switzerland) and DIG RNA Labeling Mix (Roche). Whole-mount *in situ* hybridization (WISH) was performed as described previously^3,16^. For WISH, we used plants at the anisocotyledonous stage in which GM and BM were evident.

These were immersed in fixative solution, comprising 4% w/v paraformaldehyde, 15% v/v dimethyl sulfoxide, and 0.1% v/v Tween-20 in phosphate-buffered saline (PBS), for 1 h at room temperature. DIG was detected using a DIG Detection Kit (Roche) with anti-DIG antibody (Roche).

To observe detailed spatial expression patterns, the stained samples were subsequently embedded in Technovit 7100 (Heraeus Kulzer, Wehrheim, Germany) and sliced into 10–15 μm thick sections using a HM360 rotary microtome (Thermo Fisher Scientific).

### Scanning electron microscopy

Plants were fixed in formalin-acetic acid-alcohol (FAA; 10% formalin, 5% acetic acid, 50% ethanol, v/v). The fixed samples were dehydrated through a graded ethanol series (50, 70, 90, 95, 99.5, 100%, v/v). Ethanol was subsequently replaced with a 1:1 (v/v) mixture of absolute ethanol and isoamyl acetate, followed by pure isoamyl acetate. Samples were dried in a JCPD-5 critical-point dryer (JEOL, Tokyo, Japan), sputter-coated with platinum at 20 mA for 90 s using a JEC-3000FC instrument (JEOL) in auto mode, and examined under a JSM-6510LV scanning electron microscope (JEOL).

### Sample processing and nuclei preparation

Seedlings at 21 days after sowing were dissected under a stereomicroscope. The basal region of the macrocotyledon, which encompasses both the GM and BM, was harvested, immediately frozen in liquid nitrogen, and stored at −80°C until use. Frozen tissue was carefully crushed into small pieces in liquid nitrogen using a pestle and mortar. The crushed tissue was suspended in 6 mL of precooled Honda buffer (2.5% w/v Ficoll 400, 5% w/v Dextran T40, 0.4 M sucrose, 10 mM MgCl₂, 1 mM dithiothreitol, 0.5% v/v Triton X-100, 25 mM Tris-HCl, pH 7.4) containing 1 tablet/50 mL cOmplete Protease Inhibitor Cocktail (Roche) and 0.4 U/µL RNaseOUT (Thermo Fisher Scientific). The resulting homogenate was transferred into a 10 mL tube, mixed thoroughly, incubated on ice for 5 min, filtered through a 40 µm nylon mesh cell strainer, and centrifuged at 2000 × *g* for 5 min at 4°C. The pellet was resuspended carefully in 1 mL Honda buffer and centrifuged at 2000 × *g* for 5 min at 4°C. For sorting, the nuclei pellet was resuspended in staining buffer comprising PBS containing 1 µg/mL 4’,6-diamidino-2-phenylindole (DAPI), 2% w/v bovine serum albumin, and 0.4 U/µL RNaseOUT. Nuclei were sorted by gating on the DAPI peaks and single-nucleus events using a BD FACS Aria III instrument (BD Biosciences, San Jose, CA, USA) in a small volume of staining buffer.

### snRNA-seq library construction and sequencing

The nuclei were loaded into a 10X Chromium system using a Single Cell 3′ Reagent Kit v3 according to the manufacturer’s protocol. We aimed to load ∼8000 nuclei per run. Following library construction, libraries were sequenced on an Illumina NovaSeq 6000 system or a NovaSeq X system.

### Preprocessing of raw snRNA-seq data

Raw sequencing data were mapped to the reference genome, yielding 8642 and 6711 nuclei from replicates 1 and 2, respectively, after prefiltration by Cell Ranger v7.2.0 (**Table S1**).

### snRNA-seq data analysis

Data analysis was carried out using the R package Seurat v5.0.1^94^. Data processing and plotting were performed with R/RStudio. The nuclei were filtered to remove those with fewer than 200 unique features or more than 9000 unique features. Batch effects were corrected using the anchor-based integration approach implemented in the FindIntegrationAnchors and IntegrateData functions with default parameters. Data were log-normalized using the NormalizeData function with the scaling factor set to 10,000. Variable genes were identified with FindVariableGenes using selection.method = “vst” and nfeatures = 2000. Data were then scaled with ScaleData. Principal component analysis (PCA) dimensionality reduction was performed using RunPCA for genes with high dispersion. A shared nearest-neighbor graph was constructed using the first 33 principal components with FindNeighbors, and clusters were identified using FindClusters at a resolution of 1.3. Cell clusters were visualized by uniform manifold approximation and projection (UMAP), which was performed using RunUMAP. Cluster-enriched genes were identified using FindAllMarkers with parameters logfc.threshold = 0.25 and min.pct = 0.25. GO enrichment analysis was performed based on the BLASTx best match between representative genes and *A. thaliana* protein sequences (Araport11). Orthologs were detected with OrthoFinder using the predicted protein sequences for the species and protein sequences from *A. thaliana*.

### Integration of *M. glabra* snRNA-seq and *A. thaliana* single-cell RNA-seq data

scRNA-seq data from *A. thaliana* shoot apex protoplasts^36^ were processed individually using the 10X Genomics Cell Ranger count pipeline against a reference constructed from the Araport11 reference annotations. The Seurat workflow was performed as described above. After clustering, cell clusters were annotated based on known marker genes. To identify orthologous genes for integration between *M. glabra* and *A. thaliana*, the gene with the highest expression level was selected when multiple genes were included in the same orthologous group. Integration anchors were chosen from multi-to-multi orthologs.

For cross-species single-cell integration, we utilized the reciprocal PCA (rPCA) method implemented in Seurat. Due to the evolutionary distance between the species, we chose rPCA over canonical correlation analysis (CCA) as a conservative approach to avoid overcorrection. The integration dataset was generated using 11,502 orthologous genes. Cell clusters were visualized using the methods described above.

To characterize the properties of each integrated cluster categorized into SAM cells, cluster-enriched genes were identified using FindAllMarkers with parameters logfc.threshold = 0.25 and min.pct = 0.25, among SAM-annotated clusters. To identify differences between *M. glabra* and *A. thaliana* within each integrated cluster, species-specific markers were calculated using FindConservedMarkers with the same thresholds.

---

**Figure S1.**
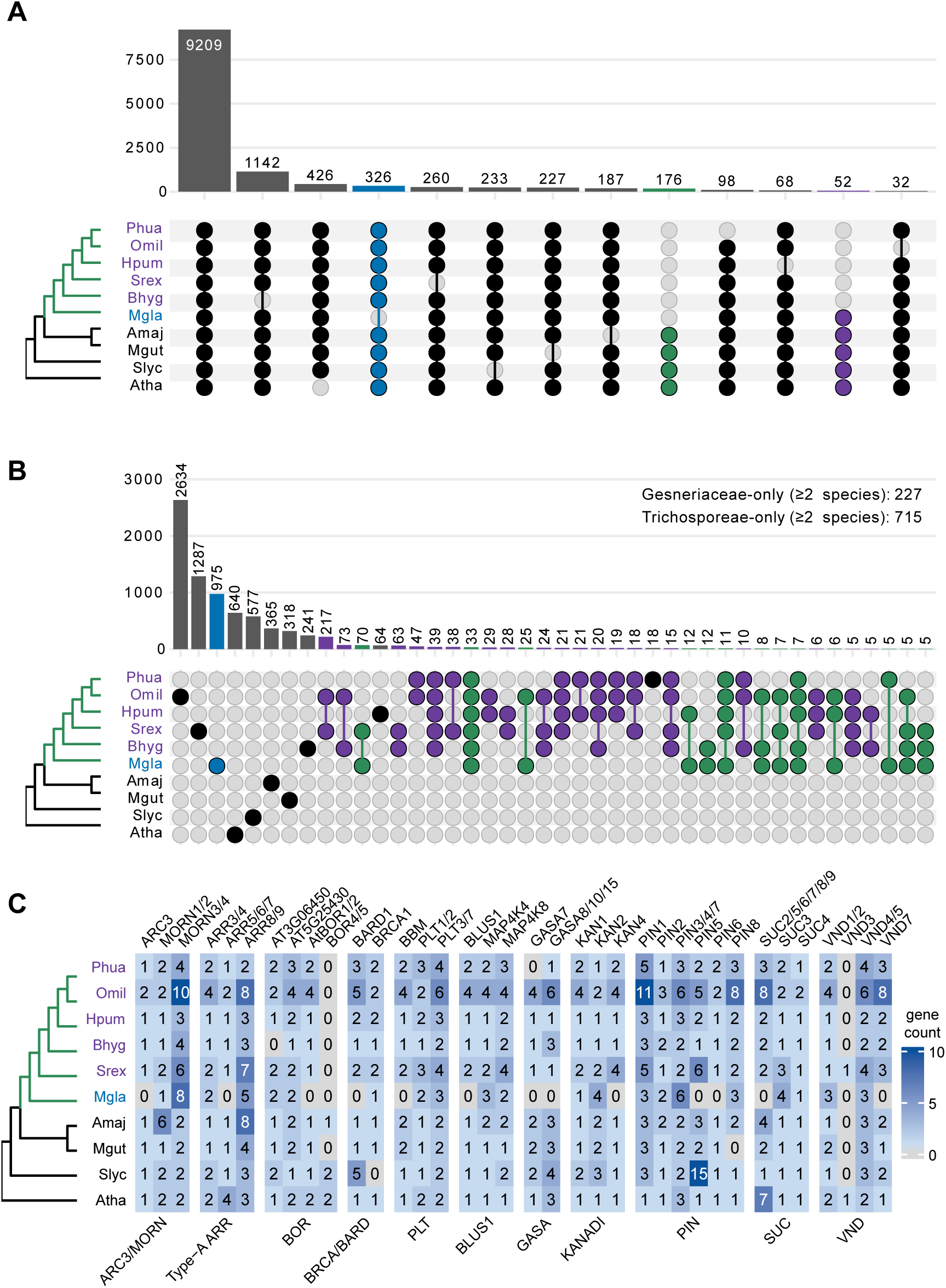
Putative losses and gains of orthologous groups in Gesneriaceae. (A, B) Number of unique and shared orthologous group (OG) losses (A) or gains (B) among the analyzed species. (C) Gene copy numbers within OGs lost in the analyzed species. Plots are color-coded to highlight specific lineages: purple indicates species belonging to the tribe Trichosporeae, and blue highlights *M. glabra*. Species abbreviations: Atha, *Arabidopsis thaliana*; Slyc, *Solanum lycopersicum*; Mgut, *Mimulus guttatus*; Amaj, *Antirrhinum majus*; Mgla, *Monophyllaea glabra*; Strex, *Streptocarpus rexii*; Bhyg, *Boea hygrometrica*; Hpum, *Henckelia pumila*; Omil, *Oreocharis mileensis*; Phua, *Primulina huaĳiensis*.

**Figure S2.**
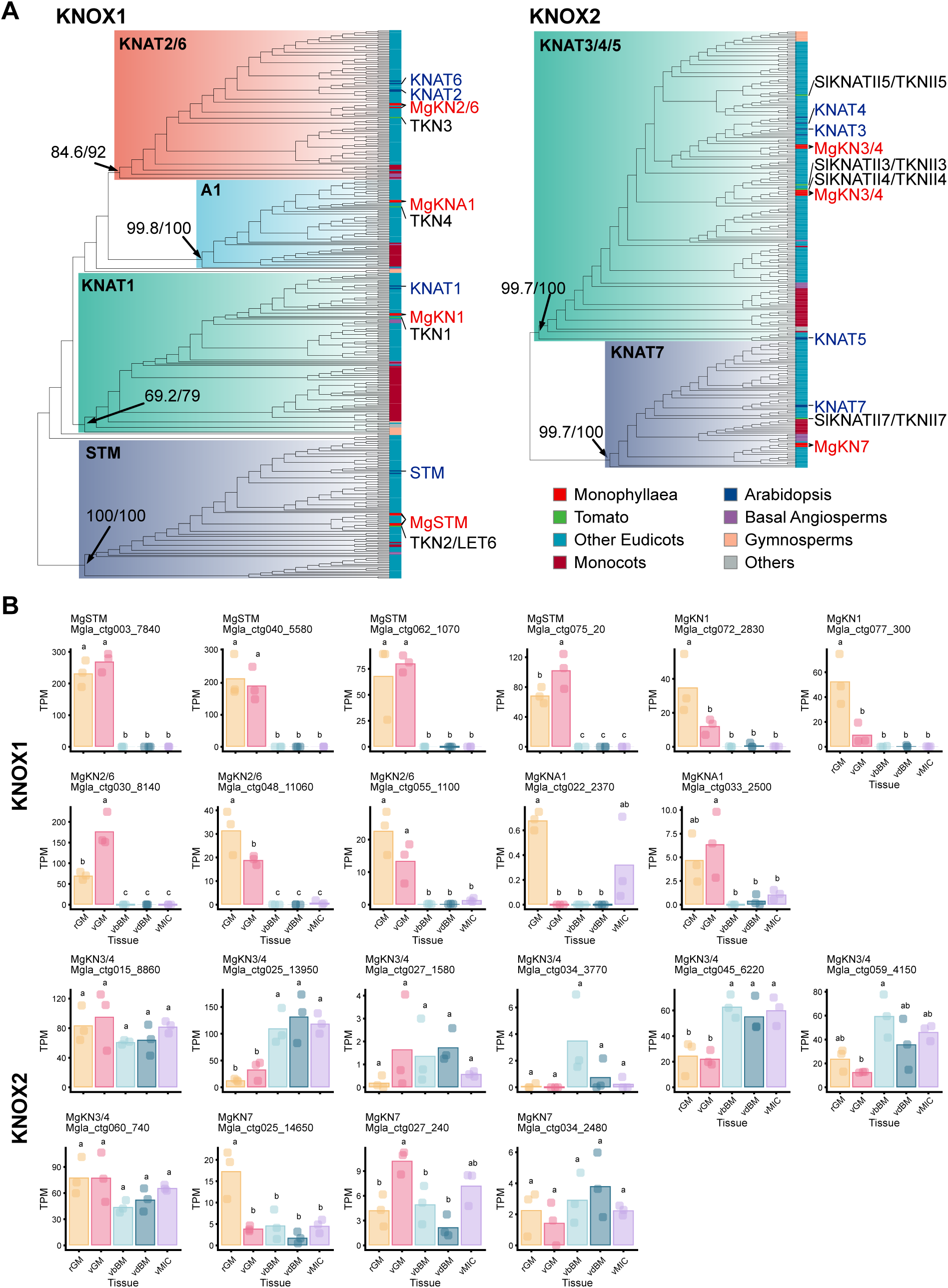
Tissue-specific RNA-seq analysis of class I and II KNOX genes in *M. glabra*. (A) Phylogenetic relationships of class I and class II KNOX (KNOX1 and KNOX2) protein sequences. Only major operational taxonomic units (OTUs) are shown. Tip colors indicate species or clades. Values at basal nodes represent Shimodaira-Hasegawa-like approximate likelihood ratio test (SH-aLRT) support (%) and ultrafast bootstrap (UFboot) support (%). (B) Expression patterns of KNOX1 and KNOX2 genes across tissues. TPM values are shown. Tukey’s HSD test, *p* < 0.001; *n* = 3. Tissues correspond to those shown in Figure 2.

**Figure S3.**
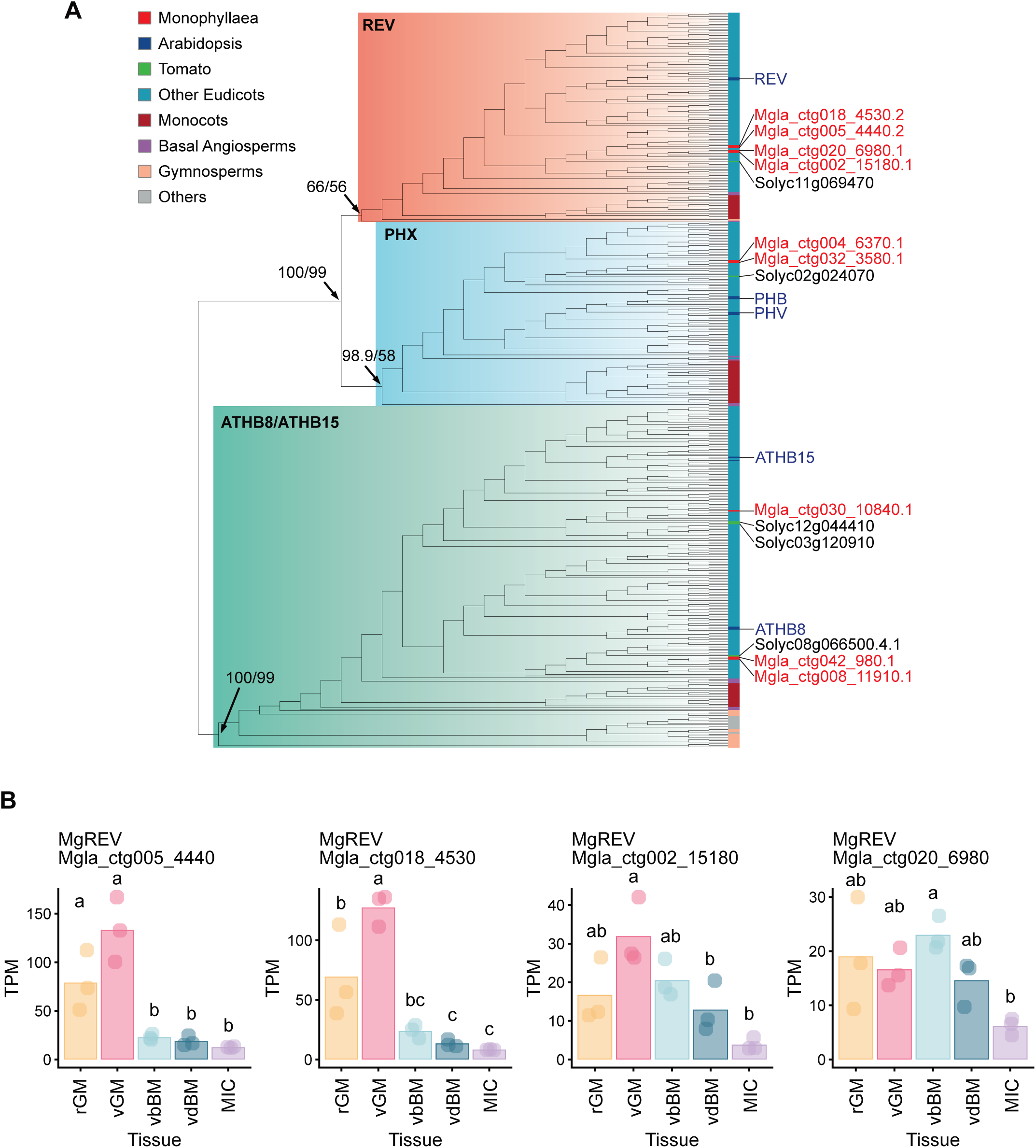
Tissue-specific RNA-seq analysis of REV genes in *M. glabra*. (A) Phylogenetic relationships of class III HD-ZIP protein sequences. OTUs are shown. Tip colors indicate species or clades. Values at basal nodes represent SH-aLRT support (%) and UFboot support (%). (B) Expression patterns of *MgREV* genes across tissues. TPM values are shown. Tukey’s HSD test, *p* < 0.001; *n* = 3. Tissues correspond to those shown in Figure 2.

**Figure S4.**
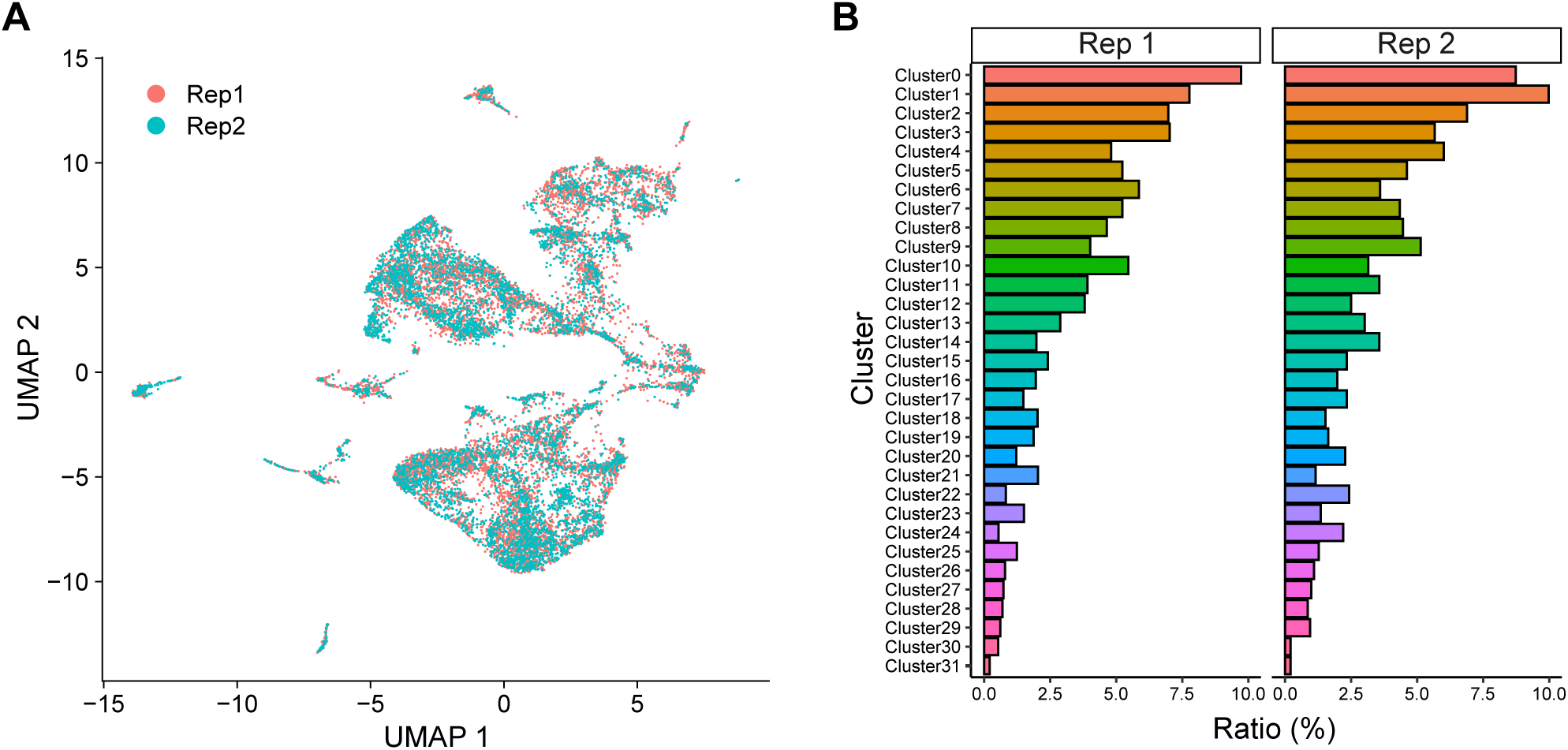
Evaluation of data reproducibility in *M. glabra* snRNA-seq. (A) UMAP visualization of the integrated snRNA-seq dataset after batch-effect correrction, color-coded by biological replicate (Rep 1 and Rep2). (B) Proportions of cells derived from each biological replicate across the individual clusters.

**Figure S5.**
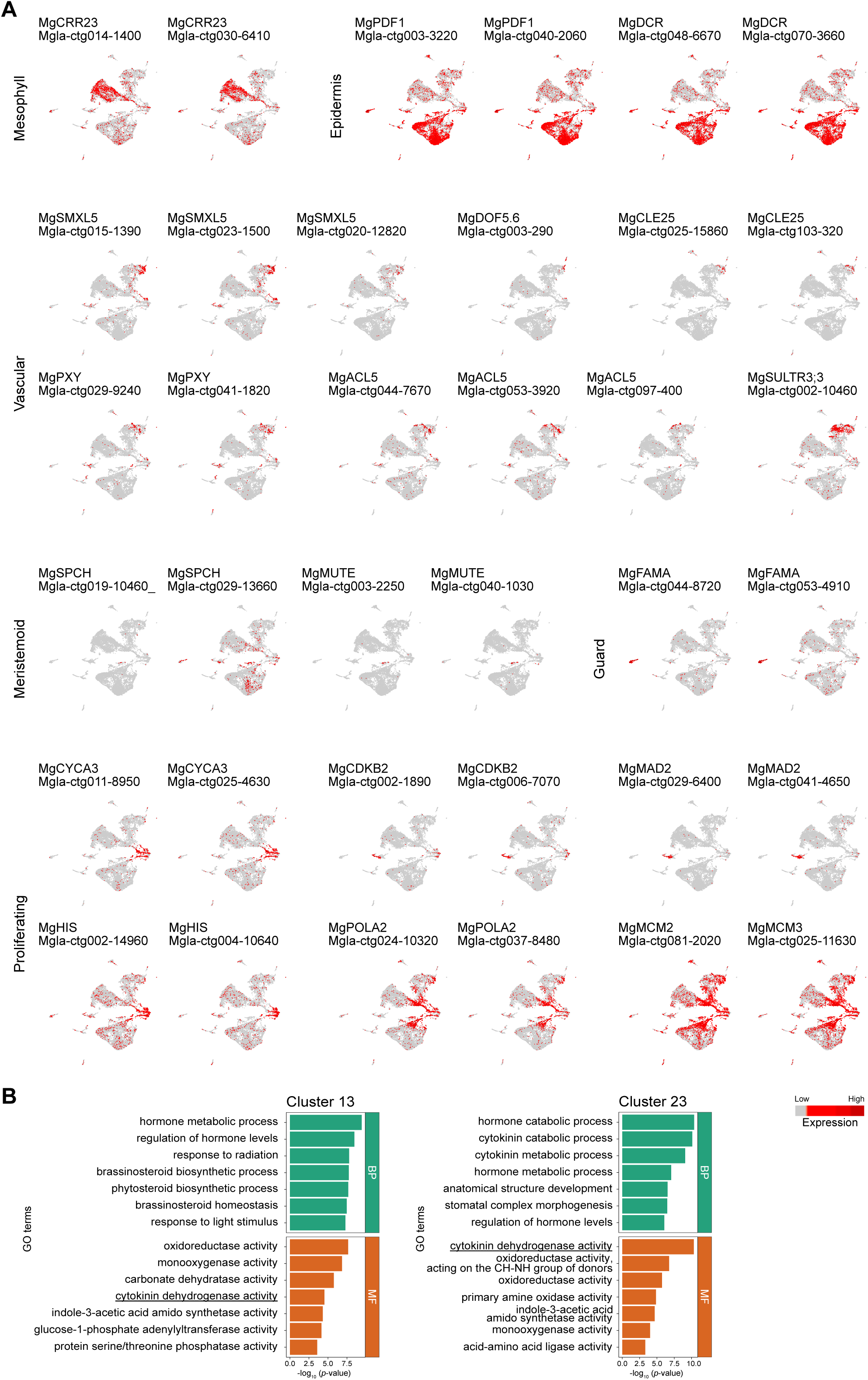
Expression of marker genes used for cell type annotation in the *M. glabra* snRNA-seq dataset. (A) UMAP plots showing expression patterns of selected marker genes for distinct cell types. (B) GO enrichment analysis of genes enriched in clusters 13 and 23. Top enriched Biological Process (BP) and Molecular Function (MF) terms are shown.

**Figure S6.**
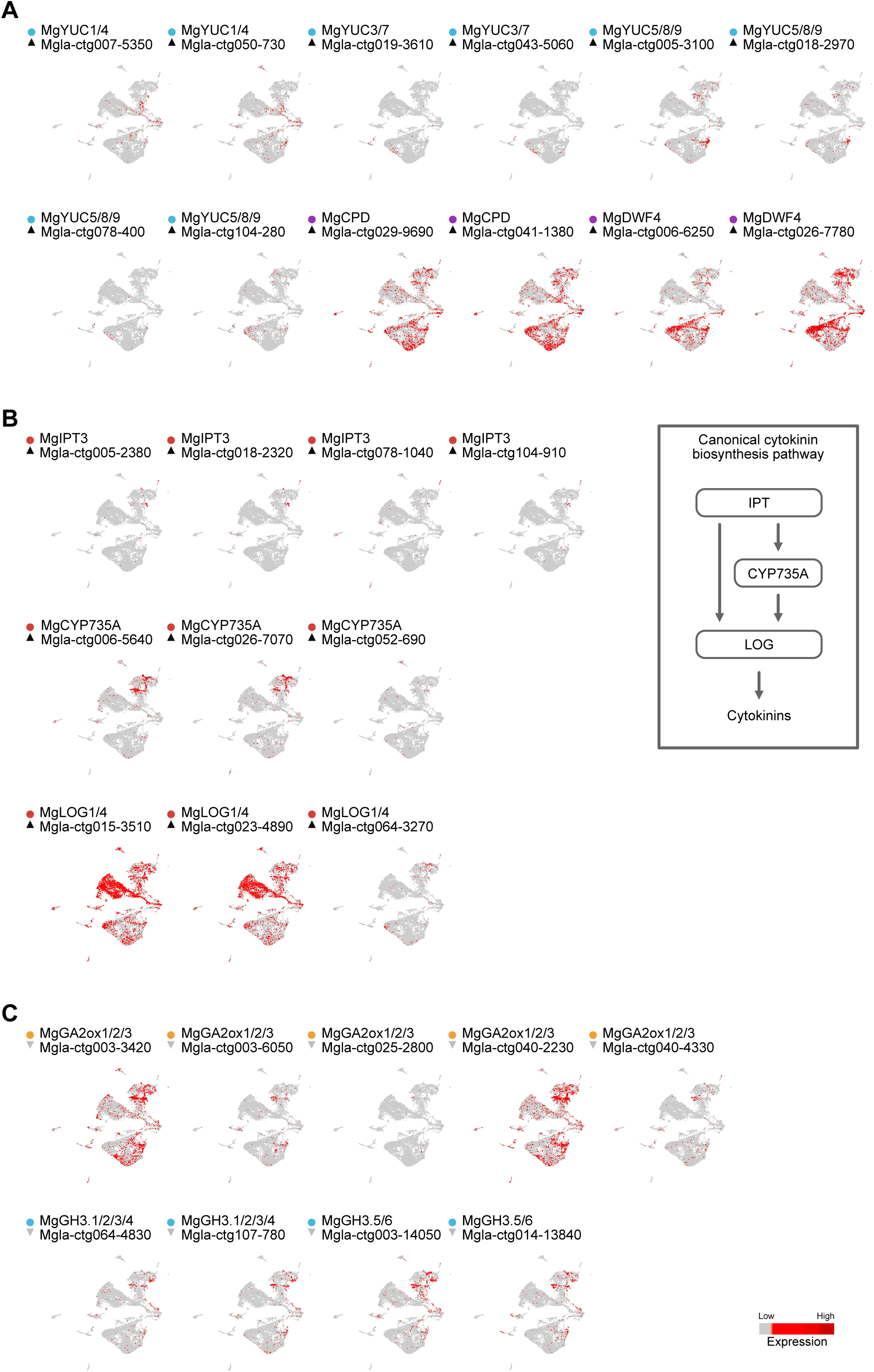
Expression profiles of phytohormone metabolic genes in meristems of *M. glabra*. (A) UMAP plots showing expression patterns of auxin and brassinosteroid biosynthesis genes expressed in BM cell populations. (B) UMAP plots showing expression patterns of cytokinin biosynthesis genes expressed in GM cell populations (left panel) and a schematic representation of the canonical cytokinin biosynthesis pathway (right panel). (C) UMAP plots showing expression patterns of gibberellin-inactivating *MgGA2ox* and auxin-inactivating *MgGH3* genes expressed in GM cell populations.

**Figure S7.**
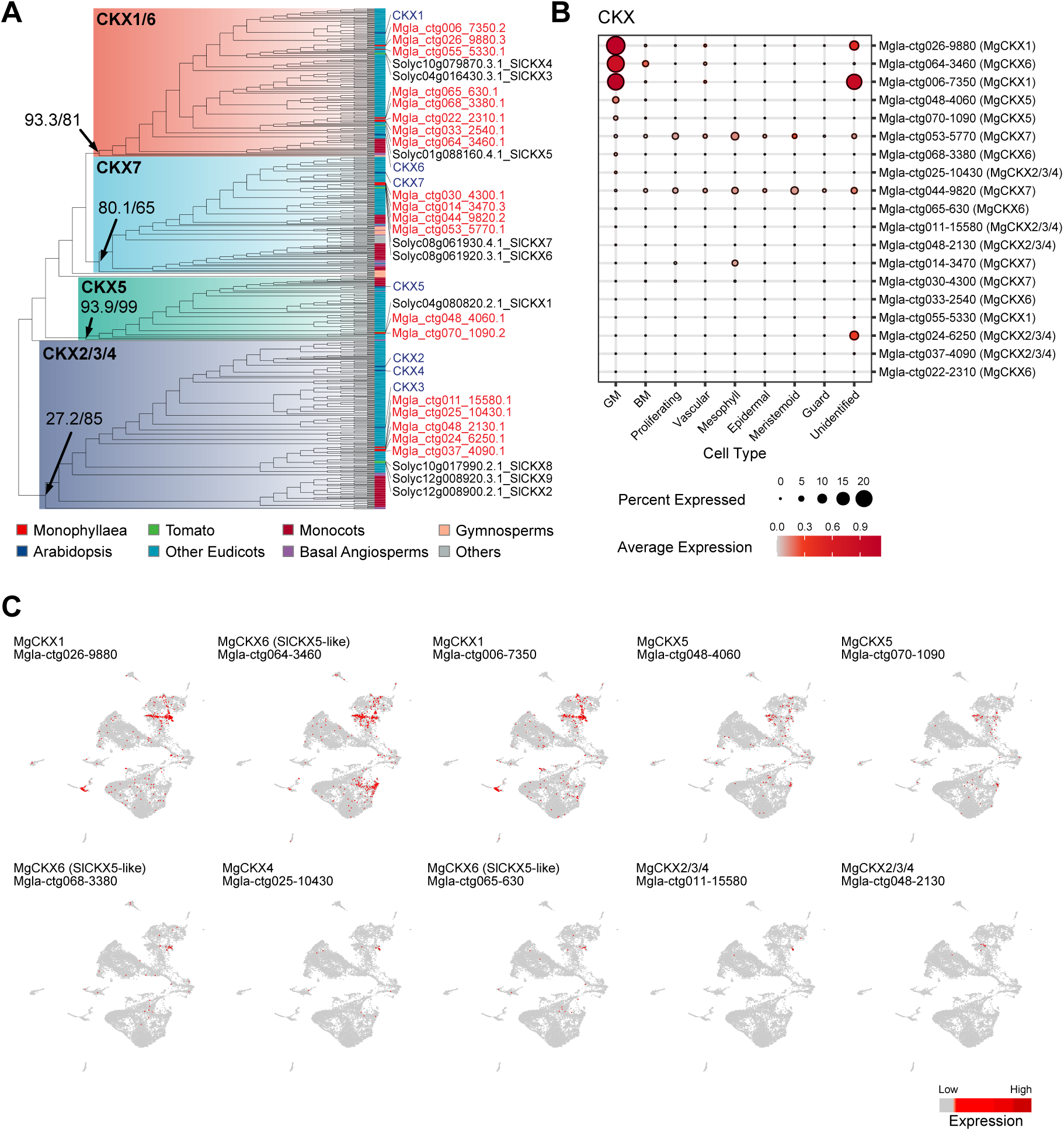
Phylogenetic and expression analyses of *CKX* genes in *M. glabra*. (A) Phylogenetic relationships of CKX protein sequences. OTUs are shown. Tip colors indicate species or clades. Values at nodes represent SH-aLRT support (%) and UFboot support (%). (B) Dot plot showing the average expression levels of *MgCKX* genes and the percentages of *MgCKX* -expressing cells across individual cell types. (C) UMAP plots for individual *MgCKX* genes expressed in the GM (Clusters 13 and 23).

**Figure S8.**
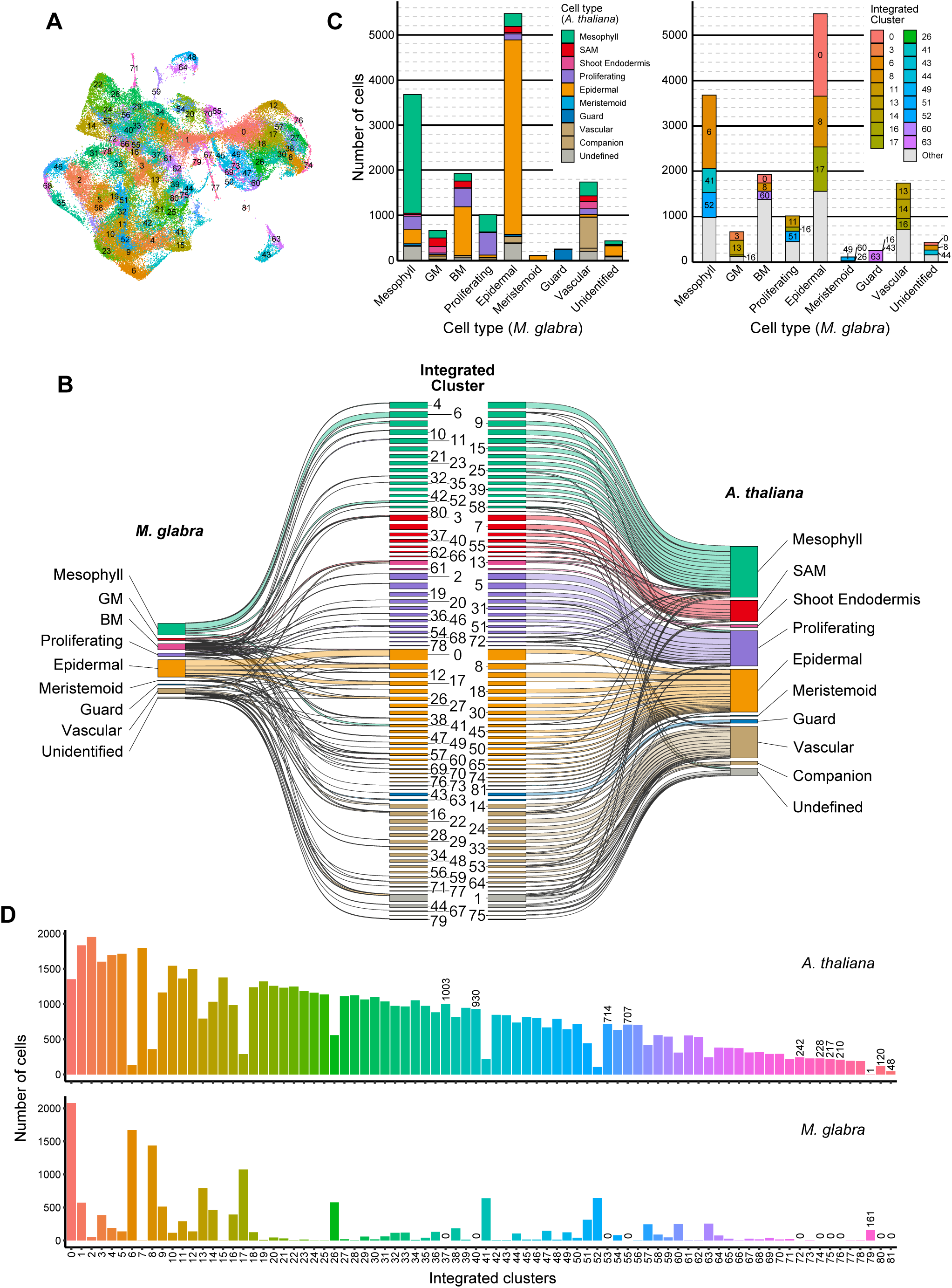
Integration and cell type correspondence of *A. thaliana* scRNA-seq and *M. glabra* snRNA-seq datasets. (A) UMAP visualization of the integrated dataset from *A. thaliana* and *M. glabra*. Cells are color-coded and labeled according to their integrated cluster numbers. (B) Sankey diagram illustrating the alignment between annotated cell types of *M. glabra* and *A. thaliana* based on integrated clusters. To minimize low-frequency background connections, links representing 1% of cells from *M. glabra* to the integrated clusters, and 10% of cells from the integrated clusters to *A. thaliana*, were excluded. (C) Proportional distribution of integrated cells for each annotated *M. glabra* cell type. The left panel displays the corresponding *A. thaliana* cell types. The right panel displays the corresponding integrated clusters, with only the top three clusters highlighted for each cell type. (D) Number of *A. thaliana* and *M. glabra* cells within each integrated cluster. Cell counts are displayed for species-biased clusters, specifically those containing zero cells from *M. glabra* or only one cell from *A. thaliana*.

